# Adaptive trade-offs between vertebrate defense and insect predation drive ant venom evolution

**DOI:** 10.1101/2024.03.06.583705

**Authors:** Axel Touchard, Samuel D. Robinson, Hadrien Lalagüe, Steven Ascoët, Arnaud Billet, Alain Dejean, Nathan J. Téné, Frédéric Petitclerc, Valérie Troispoux, Michel Treilhou, Elsa Bonnafé, Irina Vetter, Joel Vizueta, Corrie S. Moreau, Jérôme Orivel, Niklas Tysklind

**Author notes:** Corresponding author. (A.T.). These authors contributed equally to this work (Jérôme Orivel, Niklas Tysklind).

## Abstract

Stinging ants have diversified into various ecological niches, and several evolutionary drivers may have contributed to shape the composition of their venom. To comprehend the drivers underlying venom variation in ants, we selected 15 Neotropical species and recorded a range of traits, including ecology, morphology, and venom bioactivity. Principal component analysis of both morphological and venom bioactivity traits revealed that stinging ants display two functional strategies. Additionally, phylogenetic comparative analysis indicated that venom function (predatory, defensive, or both) and mandible morphology significantly correlate with venom bioactivity and amount, while pain-inducing activity trades off with insect paralysis. Further analysis of the venom biochemistry of the 15 species revealed switches between cytotoxic and neurotoxic venom compositions in some species. This study highlights the fact that ant venoms are not homogenous, and for some species, there are major shifts in venom composition associated with the diversification of venom ecological functions.

**Significance:** Venoms are under severe evolutionary pressures, exerted either on the innovation of toxins or the reduction of the metabolic cost of production (1). To reduce the metabolic costs associated with venom secretion, some venomous animals can regulate venom expenditure by metering the amount of venom injected and by switching between offensive and defensive compositions (2–2). Many ants use venom for subduing a wide range of arthropod prey, as well as for defensive purposes against invertebrates and vertebrates, but are unable to adapt venom composition to stimuli (5, 6). Consequently, the expression of venom genes directly affects the ability of ants to interact with the biotic environment, and the venom composition may be fine-tuned to the ecology of each species. A previous study showed that defensive traits in ants exhibit an evolutionary trade-off in which the presence of a sting is negatively correlated with several other defensive traits, further supporting that trade-offs in defensive traits significantly constrain trait evolution and influence species diversification in ants (7). However, the sting is not used for the same purpose depending on the ant species. Our study supports an evolutionary trade-off between the ability of venom to deter vertebrates and to paralyze insects which are correlated with different life history strategies among Formicidae.

## Introduction

Most ants secrete venom, the composition of which can vary considerably among lineages; some species have formic acid or alkaloid-based venoms, while the venoms of most stinging species are peptidic (8). The Formicidae have radiated into diverse ecological niches (9), and numerous evolutionary forces may have contributed to the shaping of their venoms.

Diet is often a potent driver of venom evolution in predatory organisms (1). Many predatory ants use their venom to capture a diversity of prey; however, several species or lineages are stenophagous (i.e. prey exclusively on a restricted group of arthropods) (10). As most stinging ants also use their mandibles to subdue their prey before delivering the paralyzing sting, mandible shape varies widely, which could be a putative driver of venom composition. The morphology of the mandibles of some predatory ants is indeed specialized to the shape of the prey (10, 11), while trap-jaw ants use spring-loaded mandibles that snap shut on prey with high speed and force (12). Foraging activities of predatory ants also range broadly from subterranean to canopy habitats, and previous research on ponerine ants suggests that arboreal constraints may influence the efficacy of venom in capturing prey (13). Other ants that live in mutualistic association with myrmecophytes (e.g. acacia ants) use their venom not for predation, but for fierce protection of the host plant (14), while in striking contrast, some stinging species lack aggression toward potential predators (15). These non-aggressive ants exhibit thanatosis (i.e. feigning death) (16), escape behavior (17), or rely on morphological attributes such as spines as a deterrent rather than using their sting against vertebrate predators (7). Among different lineages of stinging ants, venoms can exhibit very different peptide toxin profiles (18, 19), presumably in response to distinct ecological pressures. To date, no studies have integrated ecological traits, biochemical composition, and bioactivity in a phylogenetic framework to explore the factors that lead to distinct venom compositions in ants and, more broadly, very few in other venomous lineages (20).

To understand the potential evolutionary drivers underpinning stinging ant venom composition, we designed a phylogenetically nested sampling of 15 Neotropical species with contrasted ecological traits. Foraging activity (arboreal vs. terrestrial), venom function (predatory, defensive, both), mandible morphology (trap-jaw vs. normal), and prey specialization were included as ecological factors that may influence venom composition in ants. The effect of ecological traits was tested in six genera of stinging ants (i.e. *Anochetus, Daceton, Neoponera, Odontomachus, Paraponera*, and *Pseudomyrmex*). The genera *Neoponera, Odontomachus*, and *Pseudomyrmex* included both arboreal and terrestrial species to assess the effect of habitat on venom composition. *Neoponera commutata* is a known specialized termite predator (21). *Anochetus emarginatus* is also suspected to be a termite specialist (22, 23). The inclusion of *D. armigerum* allowed testing the convergent effects of trap-jaw mandibles on venom evolution with the genera *Odontomachus* and *Anochetus* (12). Within the genus *Pseudomyrmex*, we used ground-dwelling (*P. termitarius*) and arboreal (*P. gracilis*) predatory species and compared them with obligate plant-ant species (*P. viduus* and *P. penetrator*), which never use their venoms for predation (24), to examine the effects of relaxed selection pressures for predatory capacity on venom diversity. *Paraponera clavata*, notable for its painful defensive sting (25), has also been included in this panel as a venom that has evolved to effectively repel vertebrate predators. We analyzed behavior, diet, a suite of morphological traits, venom efficacy, and venom composition across phylogenetic relationships.

## Results and Discussion

### Ecological and venom-related traits

First, we collected observational data about the diet and the use of venom during prey capture or defense to fill the ecological knowledge gap for the studied species. These observations enabled us to define the ecological traits of all the species studied (**Figure 1**). We did not retain diet specialization as an ecological trait for further analysis since *A. emarginatus* appeared to be an euryphagous predator (i.e. prey on numerous classes of invertebrates) (***SI Appendix*, Figure S1)** like all other predatory species included in our study, except for *N. commutata*. We then measured how venom-related traits varied among the 15 ant species (***SI Appendix*, Figure S2, S3**, and **S4**). All the morphological data and proportions related to venom yield, venom reservoir volume, sting length, and mandibles length are presented in ***SI Appendix*, Table S1**. All studied ants use either the sting, the mandibles or both to capture prey or to defend against predators. We therefore measured the proportions of the sting and the mandibles with the hypotheses that a long sting would be associated with a defensive function, while long mandibles would allow better seizing of prey. *Pseudomyrmex penetrator* and *Pa. clavata* had the longest stings with ratios of 0.58 and 0.55 and *Odontomachus* spp., *A. emarginatus*, and *P. gracilis* the shortest (ratios ranged from 0.32 to 0.41) (***SI Appendix*, Figure S2, E, Table S1)**. *Pa. clavata* and *D. armigerum* were the species with the longest mandibles with ratios of 1.0 and 1.1, while *P. penetrator, P. viduus* and *P. gracilis* have short mandibles with an average ratio of 0.4 (***SI Appendix*, Figure S2, F, Table S1)**. We also evaluated the potency of the 15 venoms to trigger nociception in vertebrates and to paralyze and to kill invertebrate prey (***SI Appendix*, Table S2** and **Table S3**). The capacity of the venom of a given species is a product of both the venom potency and the amount of venom delivered. To be able to compare species, we therefore calculated both their nocifensive capacity (pain-inducing) and paralytic capacity by dividing the average venom yield (μg) by venom potency (μg/mL) (***SI Appendix*, Table S2 and S3**). Finally, to provide insights into the mechanism of action of the venoms, we evaluated their cytotoxicity against *Drosophila* S2 cells using two assays that measure the effect on cell metabolism and membrane cell integrity. At a concentration of 100 μg/mL, all crude venoms except those of *A. emarginatus* and *D. armigerum* were cytotoxic (***SI Appendix*, Figure S5**). The venoms of *Odontomachus* spp. were cytotoxic at high doses, affecting cell metabolism with LC_50_ values ranging from 15.8 to 42.9 μg/mL and cell membranes with LC_50_ values ranging from 18.5 to 49.0 μg/mL. The venoms of *Neoponera* spp. were more cytotoxic with LC_50_ ranging from 5.5 to 8.3 μg/mL and from 5.8 to 14.8 μg/mL for cell metabolism and cell membrane integrity assays, respectively. The *Pseudomyrmex* spp. venoms were very cytotoxic, and the venoms of *P. penetrator* and *P. termitarius* were the most potent, impacting both cell metabolism (LC_50_ of 0.05 and 0.24 μg/mL for *P. penetrator* and *P. termitarius*) and cell membrane integrity (LC_50_ of 0.08 and 0.23 μg/mL for *P. penetrator* and *P. termitarius*). *Paraponera clavata* venom was cytotoxic but was more potent on cell metabolism (LC_50_ of 2.72 μg/mL) than on cell membrane integrity (LC_50_ of 31.03 μg/mL). Detailed results are presented in ***SI Appendix*, Table S4** and **Figure S5**. Since no cytotoxic activity was observed for *A. emarginatus* and *D. armigerum* crude venoms, we tested the effects of these venoms on cell membrane potential on S2 cells. A significant decrease in KCl-induced membrane depolarization was observed after incubation with both venoms, indicating an inhibition effect on ionic conductance ***SI Appendix*, Figure S6**.

**Figure 1.**
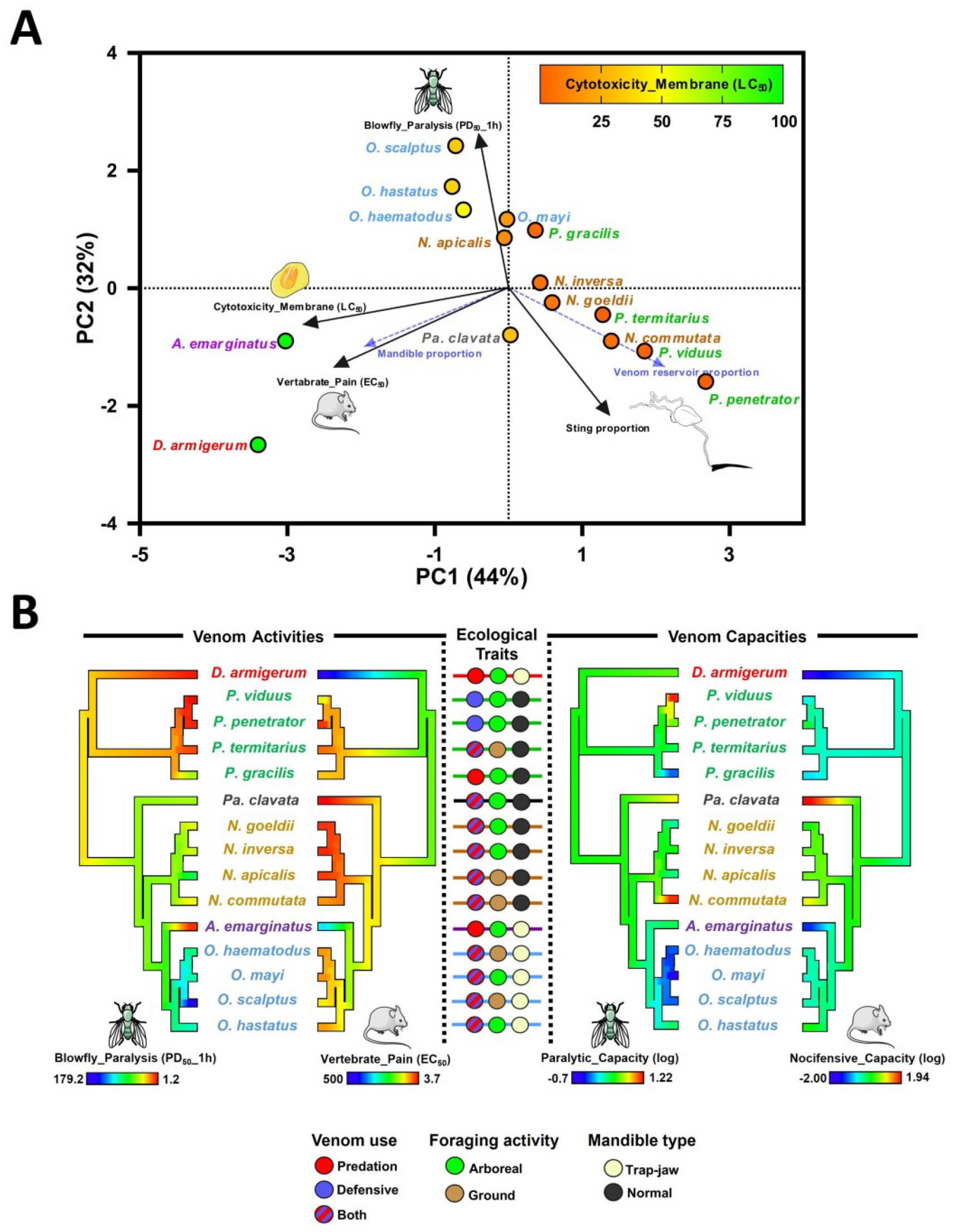
Venom bioactivity and morphological traits in 15 ant species. **A)** Principal component analysis of 15 ant species defined by venom bioactivities and morphological features. PCA revealed two functional strategies among species based on the cytotoxicity of venom. The significance of each PC axis, and of loading of each trait have been tested by the PCAtest R package (26). Traits having significant loadings on PC 1 and PC 2 are represented with black arrows, while others are represented with dashed blue arrows. Plot points are colored in a gradient based on membrane cytotoxicity values (LD_50_) as indicated by the scale bar at the top right. See also ***SI Appendix*, Figure S7** for PCA on the dataset featuring all traits. **B)** Ancestral state reconstructions of the insect paralytic activity (PD_50__1h), vertebrate pain activity (EC_50_), paralytic capacity, and nocifensive capacity of crude ant venoms, estimated by using the Phytools R package (27). Since *D. armigerum* venom was inactive on F11 cells, we used an arbitrary high value of 500 μg/mL for vertebrate pain (EC_50_). Capacities were calculated by dividing the average venom yield (μg) by the venom potency to paralyze blowfly (PD_50__1h, μg/g) and to cause pain (EC_50_, μg/mL). Venom capacities have been log-transformed. The scale bar indicates trait values from low (cool colors) to high potencies (warm colors) for venom activities and from low (cool colors) to high venom capacities (warm colors).The phylogenetic tree was reconstructed by using transcript sequences of 566 BUSCO genes expressed in the body of ant species.

### Venom-related traits reveals the evolution of two functional strategies

A principal component analysis (PCA) was done on the dataset featuring the venom activities, cytotoxicity, and morphological traits. The first two axes of the PCA accounted for 76% of the total variation, with axes 1 and 2 explaining 44% and 32% of the total variation, respectively (**Figure 1A**). The vertebrate pain activity and insect cell cytotoxicity have significant loading on PC1 while prey paralysis and sting proportion have significant loading on PC2. PCA revealed that two different venom strategies are used by the studied ant species. The strategy used by *A. emarginatus* and *D. armigerum* can be defined as species with a non-cytotoxic venom that is capable of paralyzing blowflies efficiently but has a poor capacity to induce pain in vertebrates. All other species with the second strategy are distributed along a venom cytotoxicity gradient, with the most cytotoxic venoms causing more pain in vertebrates, paralysis and lethality in flies, and tending to have a longer sting.

Among ponerine species, we noted a major shift in insect paralytic activity with *A. emarginatus*, whose venom is 13 to 22 times more paralytic than those of *Odontomachus* species and at least 5 times more paralytic than those of *Neoponera* species (**Figure 1B**). Reconstruction of the ancestral state of vertebrate pain activity illustrates how the venoms of *D. armigerum* and *A. emarginatus* lack vertebrate pain-inducing ability (e.g. the venom of *A. emarginatus* is 3 to 5 times less active on vertebrate sensory neurons than that of *Odontomachus* species and 100 times less than that of *Pa. clavata*). The amount of venom varies greatly among species, which greatly impacts their paralytic and nocifensive capacities. It is therefore worth noting that the venom of *A. emarginatus* and *D. armigerum* has a very low nocifensive capacity, whereas the venom of *Pa. clavata* and to a lesser extent *N. commutata*, has a high nocifensive capacity (**Figure 1B**).

Altogether, the venom activities and capacities align well with the lifestyle and the venom use of each species. *Anochetus emarginatus* rarely stings defensively, and the sting is not painful, causing only a slight itch (personal observation A.T.) which may explain the cryptic lifestyle of most *Anochetus* species (28). When disturbed, *A. emarginatus* primarily utilizes its trap-jaw mandibles to bite and bounce off intruders (personal observation A.T. and (29)). *Daceton armigerum* does not sting defensively (personal observation A.T.) and we showed that the crude venom caused no pain-inducing activity. To avoid predation, *D. armigerum* has a very thick cuticle covered with thoracic spines and has adopted an arboreal lifestyle, living in polydomous nests sheltered in hollow branches (30, 31). The defensive constraint against vertebrates in *Pa. clavata* and *N. commutata* may be more pronounced than in other species, since they ranked first and second in nocifensive capacity. Because of the large size of workers, they nest directly in the ground, making the colony attractive in terms of nutritional resources and highly vulnerable to vertebrate predation. Among ponerine ants, the venom of *Odontomachus* spp. has a low paralytic activity and capacity. *Odontomachus* spp. capture their prey with trap-jaw mandibles and do not always use their venom, depending on the prey type (32). In the *Pseudomyrmex* clade, the venom of the plant-ant species (i.e. *P. penetrator* and *P. viduus*) has a very high paralytic capacity on insects, in marked contrast to that of *P. gracilis*. Although *P. viduus* and *P. penetrator* species do not use their venom for predation, they are subject to strong selective pressures to defend host plants against both grazing insects and vertebrates. *Pseudomyrmex penetrator* venom is also highly effective at inducing pain in vertebrates (***SI Appendix*, Tables S2**).

### Correlation among traits

Comparative phylogenetic generalized least squares (PGLS) regression revealed several significant correlations among traits (**Figure 2**). For the evolutionary impact of ecological traits, we found that both venom use and mandible type significantly correlate with venom bioactivities and morphological traits, while there is no correlation between foraging activity (arboreal vs. terrestrial-foraging species) and any other traits (**Figure 2A**). We show that the metabolic cost of toxin secretion is reduced in stinging ant species with relaxed selective pressure for defensive function, since they produce less venom than other ants (proportion of venom reservoir volume (*P* = 0.003)). The defensive function significantly affects the properties of venom: the venoms of species that use their venom defensively generally have greater cytotoxicity (cytotoxicity_metabolism, *P* = 0.021; cytotoxicity_membrane, *P* = 0.031), as well as greater vertebrate pain activity (*P* = 0.03), associated with higher nocifensive capacity (*P* = 0.03) than species that use venom exclusively for predatory purposes. Counterintuitively, the predatory use of venom significantly reduces the potency against prey, measured as paralysis (*P* = 0.024) and lethality (P = 0.007) in blowflies, suggesting that some predatory species may compensate low venom activity to capture prey with other adaptations such as trap-jaw mandibles. Mandible strike performances vary among trap-jaw species and may however have a variable influence on the venom activity (33). In this study, the presence of trap-jaw mandibles has no effect on venom activity against blowflies, both paralysis and lethality, but was correlated with low vertebrate pain activity (*P* = 0.016), low venom volume (*P* = 0.022), a smaller sting (*P* = 0.010), low nocifensive capacity (*P* = 0.021), and low cytotoxicity (cytotoxicity_metabolism, *P* = 0.007; cytotoxicity_membrane, *P* = 0.036). The presence of specialized mandibles is therefore associated with an overall decrease in the defensive function of venom.

**Figure 2.**
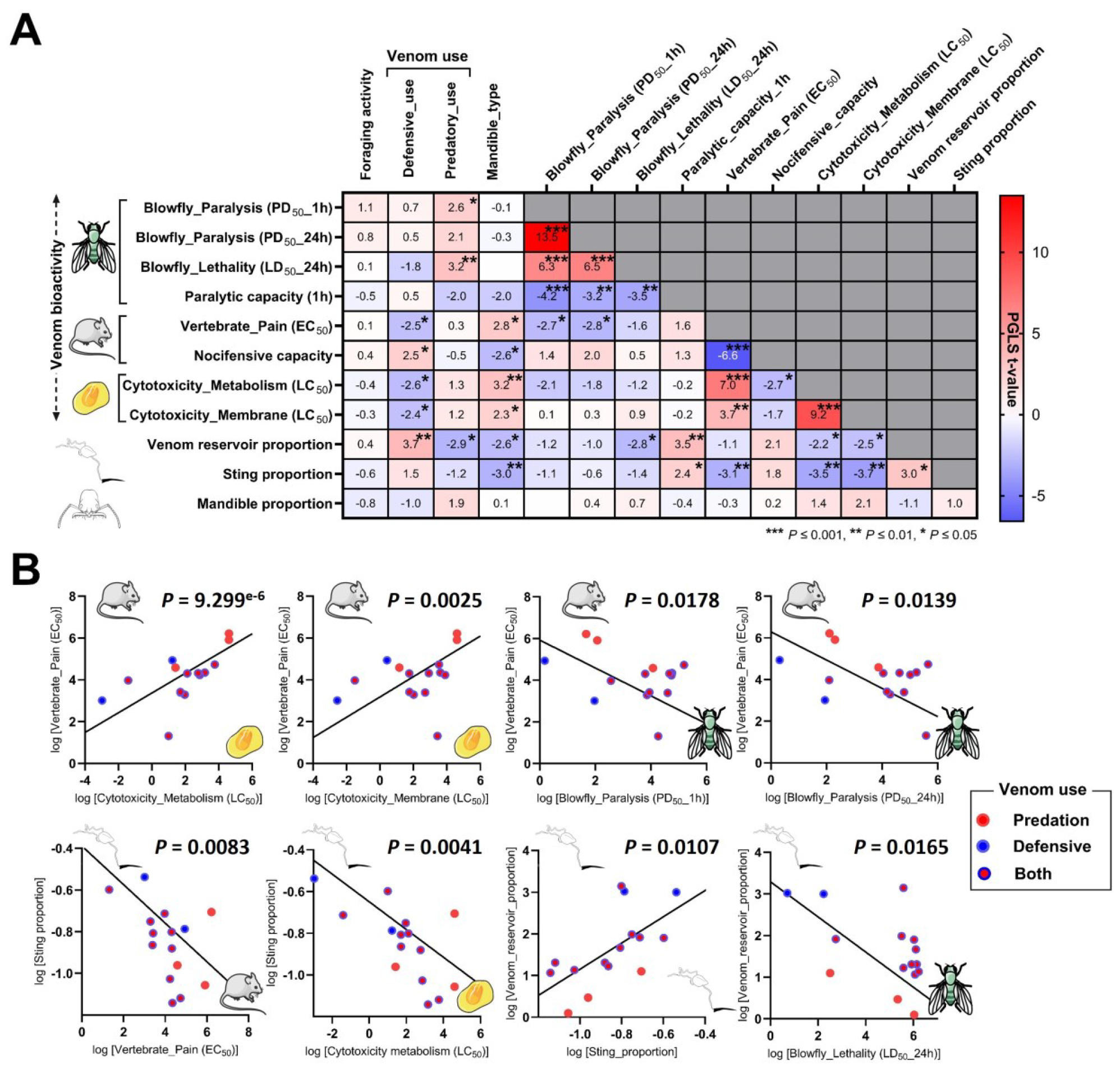
Comparative phylogenetic analysis. **A)** Phylogenetic generalized least squares (PGLS) analysis among ecological traits, venom bioactivity, and morphological traits in the 15 ant species. As “venom use” is a multi-state discrete variable with non-ordinal properties that contain a category “both”, we decomposed that trait into two binary discrete variables (defensive_use and predatory_use) having only two states (yes or no). Heatmap with PGLS t-values and statistical significance. Positively correlated values are in red and negatively correlated values are in blue. **B)** PGLS linear regressions of several significantly correlated traits.

The data showed that the longer the sting, the more pain activity (*P* = 0.008) and cytotoxicity (cytotoxicity_metabolism, *P* = 0.004; cytotoxicity_membrane, *P* = 0.002) were found in the venom, which is consistent with an anti-vertebrate role for a long sting. PLGS regression showed a significant positive correlation between the venom reservoir proportion with both the sting proportion and lethality in blowflies (*P* = 0.011). Pain activity in vertebrates is strongly positively correlated with the cytotoxicity of venoms (cytotoxicity_metabolism, *P* < 0.001; cytotoxicity_membrane, *P* = 0.002) but negatively correlated with the paralysis in blowflies (blowfly_paralysis_1h, *P* = 0.018; blowfly_paralysis_24h, *P* = 0.014) (**Figure 2B**). This is suggestive of a trade-off between vertebrate pain-inducing activity and insect-predation activity in these ant venoms. Such a trade-off might translate to life history strategies, where species with potent paralytic venom against prey have reduced capacity to deter vertebrate predators and are therefore prone to adopt alternative defensive strategies, such as cryptic habits, nesting strategies (e.g. polydomous nest) or promoting behavioral (e.g. thanatosis or escape behavior) and morphological (e.g. thick cuticle and spines; body size reduction) anti-predation co-adaptations.

### The venom composition of stinging ants

To understand the biochemical mechanisms underlying the observed variations in venom efficacy, we examined the venom composition of each of the 15 species (**Figure 3)**. For further details on venom composition, see ***SI Appendix*, Figures S8-S19** and **Dataset S1**. Overall, our investigations revealed that the venoms displayed heterogeneity in composition, with a considerable turnover of peptide families across genera and without any correlation with the ecological traits considered. Most of the peptide families were genus specific with only family 9 (ponericin G) shared between *Odontomachus* and *Neoponera* venoms. The venoms of the three species that were unique for their genus (i.e. *Pa. clavata, A. emarginatus*, and *D. armigerum*) showed a very distinctive profile dominated by families of neurotoxins not shared with any other venom. Our results also showed that many venom peptide families are shared by species of the same genus, but the proportion varied.

**Figure 3.**
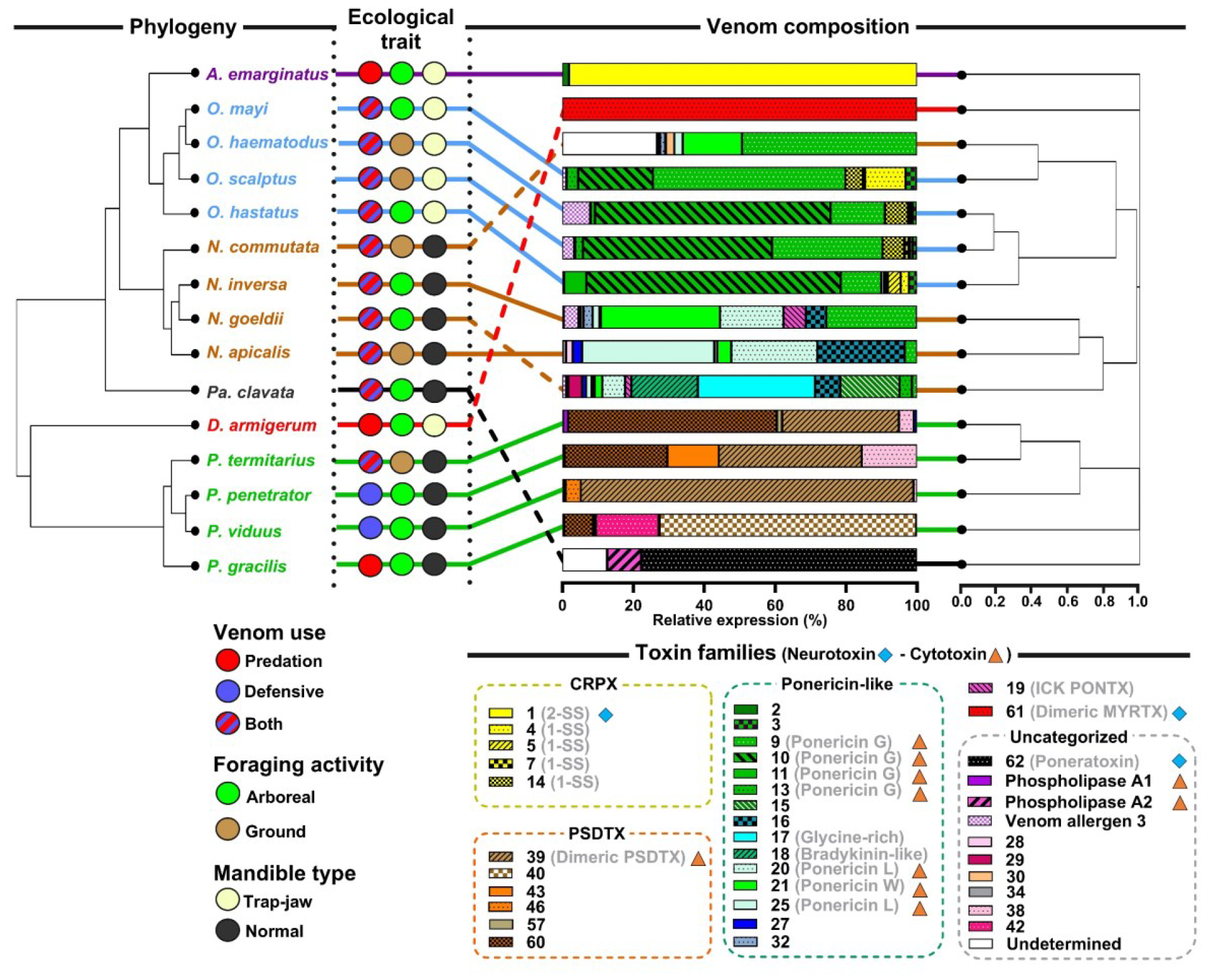
Comparison of venom composition and phylogenetic relationships among 15 ant species. Venom composition cladogram is based on the relative expression (TMM) of transcripts identified as toxins in each venom gland transcriptome and converted into Bray-Curtis distance matrix and hierarchical cluster analysis was performed by using the complete linkage method of hclust() function with the R software. Only families with a relative expression value >1% in at least one species are shown in the color key. Toxin families are grouped by precursor clades (***SI Appendix*, Figure S19**). Blue diamonds and orange triangles indicate neurotoxic and cytotoxic peptide families, respectively, based on the literature. In contrast to the other species, *A. emarginatus, D. armigerum*, and *Pa. clavata* have convergently evolved a venom composition dominated by neurotoxic peptides.

Among the species that use their venom exclusively for predation, *A. emarginatus* and *D. armigerum* are associated with a complete shift in venom bioactivities that correlate with a switch to a neurotoxic venom composition. The venom of *D. armigerum* showed a unique profile, as previously reported (18), and our analysis confirmed that this venom consists of a single family of peptides (dimeric MYRTX, family 61) that display some amino acid sequence similarity with the neurotoxic U_11_ venom peptide from *Tetramorium bicarinatum* (34) (***SI Appendix*, Figure S20**). Given the lack of cytotoxicity but strong paralytic activity on blowflies and inhibition of cell membrane potential, these data suggest that *D. armigerum* has an insect neurotoxic venom. In addition, *A. emarginatus* has a non-cytotoxic venom that is much more paralytic to insects than the other ponerine venoms studied. Given the dominance of family 1 peptides in *A. emarginatus* venom (98% of relative expression), it is likely that 2-SS CRPXs (family 1) are responsible for most, if not all, of the paralytic effects upon prey. Since one of the peptides from family 1 (i.e. Ae1a) has shown inhibition (at a high concentration) on the human voltage-gated calcium channel (Ca_V_1), it is possible that 2-SS CRPXs are neurotoxins (35). Although the venoms of *D. armigerum* and *A. emarginatus* share similar non-cytotoxic, insect neurotoxic activity, the difference in composition suggests that they have evolved independently. By contrast, the stings of other ponerine *Odontomachus* spp., and *Neoponera* spp. induce sharp pain (36, 37) and showed stronger pain activity in our assay. Ponericins are very prevalent in the venom composition of *Odontomachus, Neoponera*, and several other ponerine ants (37–37). Ponericins are multifunctional cytotoxic peptides acting on the cell membranes (40, 41), and those from the venoms of *N. apicalis* and *N. commutata* are known to cause pain in mammals and to paralyze insects (42). There is compelling evidence that membrane-active venom peptides contribute to the defensive role of multiple venomous arthropods against vertebrates (5, 43–43).

Neurotoxic peptides are not exclusive to ants that rely on venom solely for predation. The venom of *Pa. clavata* is largely dominated by poneratoxin (family 62) (46), a pain-inducing neurotoxin that efficiently modulates vertebrate voltage-gated sodium (Na_V_) channels while paralyzing insects only at very high doses (47). We showed that the venom of *Pa. clavata* also exhibits cytotoxicity, which is likely attributed to phospholipase A_2_ (PLA_2_), present in this venom at higher levels than in other ants. Since *Pa. clavata* also uses its venom for predation, cytotoxicity may also be a means of subduing arthropod prey. Alternatively, cytotoxicity may also be a means of maintaining a multifunctional defense against predators. In this way, the venom retains a general repellent effect that ensures a baseline defense of the colony in a scenario where a predator would acquire resistance to neurotoxins. This hypothesis is supported by a previous study of the venom of the seed-harvesting ant *Pogonomyrmex*, which uses its venom primarily for defense against vertebrates, and has evolved venoms dominated by vertebrate-selective peptides that target Na_V_ channels, but still contain a peptide that is cytotoxic to vertebrate cells (48).

Our results indicated that the four *Pseudomyrmex* species have highly cytolytic venoms and different venom profiles to other ant species, with peptide families not shared with the other species studied. We found that *P. gracilis* shares few similarities with the other three *Pseudomyrmex* species, and that the different venom families are present in different proportions among the species. Among these families, family 39 corresponded to the myrmexins (renamed here the dimeric pseudomyrmeciitoxin (PSDTX)), a group of dimeric peptides first described in the venom of *P. triplarinus* (49) which are highly cytotoxic to insect cells (50). Dimeric PSDTXs largely dominate the venom of *P. viduus* (94% of relative venom expression), which was found to be the most paralytic and lethal venom on blowflies. The other families consist of cysteine-free polycationic peptides which may also contribute to the observed cytotoxicity. The venom of *P. penetrator* is at least 3 times more cytotoxic than in other *Pseudomyrmex*, at least 72 times more cytotoxic than *Neoponera*, and at least 231 times more cytotoxic than *Odontomachus* (***SI Appendix*, Table S4**). This is associated with high efficacy for inducing pain in vertebrates and paralyzing insects. This example highlights the fact that the trade-off between vertebrate and insect-predatory venom activity may be disrupted by very high cytotoxicity.

## Conclusion

Here we use a multi-pronged approach to test foraging activity, venom function, and mandible morphology as evolutionary drivers underlying venom variation in ants. We show that ant venoms can be highly heterogeneous and that there have been major shifts in venom composition and bioactivities, even among phylogenetically close species. The ecological role of the venom, offensive or defensive, and even more so the breadth of biological targets, is arguably the dominant constraint in the evolution of stinging ant venom cocktails. Metering the amount of venom produced is the swiftest way to adapt to different ecological constraints (51) such as for *N. commutata*, which has a quite similar venom composition profile to other congeneric ponerine species but produces large amounts of venom. Evolution may further fine-tune venom composition toward ecologically relevant cocktails exhibiting different functional strategies based on either neurotoxins or cytotoxins. Cytotoxic toxins, which do not require a specific pharmacological receptor, are likely to offer an evolutionary advantage to species that need to target a wide range of both vertebrate and arthropod organisms with highly divergent nervous systems. In some lineages of ants, however, evolution may have favored neurotoxin-based venoms as the range of biological targets narrowed. A reasonable explanation for the prevalence of neurotoxic-based venoms in ants would be that cytotoxic peptides often act at high concentrations compared to neurotoxins (47), and are therefore likely to be associated with higher metabolic costs. Toxin innovation in the venoms of the Formicidae may therefore facilitate diversification of lifestyle and morphology, and ultimately contribute to speciation.

## Material and Methods

### Ants and venom samples preparation

Live specimens of worker ants from different colonies for each species were collected in French Guiana. To identify genes involved in venom production, we aimed to generate whole-body (i.e. head and thorax) and venom gland transcriptomes for each species, and then, subtract genes expressed in the whole-body transcriptome from those in the venom gland, to identify those genes expressed largely in venom glands. For each species we dissected: 1) both venom glands and venom reservoirs from 100 live workers per species in ultrapure water and immediately placed in 1 mL of RNAlater™ (Thermo Fisher Scientific, Waltham, MA, USA); and 2) the head and thorax of 2-3 workers in 1 mL of RNAlater™. Samples were stored at –80°C prior to RNA extraction. Crude venom samples were prepared by dissecting ant venom reservoirs in ultrapure water, then pooled in 10% acetonitrile (ACN) in ultrapure water and stored at –20°C prior to freeze-drying. Venom samples were then loaded onto a 0.45 μm Costar^®^ Spin-X tube filter (Corning Incorporated, Corning, NY, USA) and centrifuged at 12,000 g for 3 min to remove tissues from the venom apparatus. Filtered venom samples were then lyophilized, accurately weighed, and stored at –20°C until further use.

### Transcriptomics

The RNAlater™ was removed from the tubes containing the samples using a glass Pasteur pipette. Subsequently, the glands or ant bodies (head and thorax) were then disrupted using a TissueLyser II (Qiagen, Germantown, MD, USA) in RLT buffer containing 10% (v/v) of 2-mercaptoethanol (Rneasy Mini Kit, Qiagen). RNA was isolated using a phenol-chloroform (5:1) solution, followed by washing with chloroform-isoamyl alcohol (25:1) to remove any phenol traces. The RNA was bound to a Qiagen column and washed according to the manufacturer’s instructions. DNAse I (Roche Diagnostics GmbH, Mannheim, Germany) was added to eliminate any remaining DNA fragments. The RNA was eluted using sterile water, and total RNA was quantified using a Qubit 3.0 fluorometer (Thermo Fisher Scientific, Waltham, MA, USA) with the RNA HS assay kit (Life Technologies Corp., Carlsbad, CA, USA). A NanoDrop 2000 UV-Vis spectrophotometer (Thermo Fisher Scientific) was used to determine the 260/280 and 260/230 nm ratios. Finally, the purified RNA was treated with RNAstable™ LD (Biomatrica, San Diego, CA, USA) and dried using a Speed Vac (RC1010, Jouan, Saint Herblain, France) and sent for sequencing.

### Sequencing

Samples were sent to IGA Technologies Services (Udine, Italy) where samples were suspended and RIN number checked before preparation of the mRNA-seq stranded libraries. All libraries were pooled and sequenced with RAPID 2 × 250 bp runs on a HiSeq2500 Illumina sequencer (480 Million reads). Raw sequences were demultiplexed and used in downstream analyses. The resulting library size ranged from 18.4 M to 28.3 M reads and were 22.1 M reads on average.

### Transcriptome assembly, annotation, and quantification of gene expression

Recent RNA-Seq advancements offer cost-effective transcriptome data collection, yet reconstructing full-length transcripts in non-model species from short reads remains challenging. A cautious approach consists of testing different programs and selecting the one producing the best expected de novo assembly (52). Three commonly used programs were tested: SOAPdenovo-Trans v1.03 (53), rnaSPAdes v3.13.1 (54) and Trinity v2.6.6 (55). Before assembly, the libraries were trimmed with cutadapt v2.3 (56). For the pilot run, the assemblies with the three programs were made on two species, *Anochetus emarginatus* and *Pseudomyrmex gracilis*. For the latter species, a genome and transcriptome were available (57). The resulting assemblies were compared with TransRate v1.01 (58), BUSCO v3.0.1 and the hymenoptera_odb9 database (59), and for *P. gracilis* only, with RNAQuast v2.1 (60) which computes some metrics with the use of a genome of reference. Based on the results and metrics obtained with the last three tools (data available on request), Trinity assembler was chosen for assembling the remaining 12 species using both venom glands and bodies libraries.

Homologies with known proteins were searched with Blastx on the curated UniProt-SwissProt database (accessed the 1st September 2019) and ToxProt database. Open reading frames (ORFs) were searched with TransDecoder (61). The minimal length of the ORF was set to 10 amino acids in order to keep the potential small venom peptides. Homologies of the predicted ORFs with proteins from the UniProt-SwissProt database were searched with BLASTp. Additional protein information was searched through the pfam database with hmmscan. The presence of rRNA was searched with Barrnap. The presence signal peptide was searched with SignalP v4.1 (61, 62). The transmembrane site was searched with Tmhmm v2.0 (63). Transdecoder was run again to integrate the BlastP and Pfam criteria in the coding region selection. Following the Trinity procedure, venom glands and bodies libraries were separately mapped and quantified using Bowtie2 (63, 64) and RSEM (65) using TMM normalization method. All the annotation results were integrated in a sqlite database with Trinotate v3.2 (66).

### Proteomics

LC–MS profiling of the crude venoms was carried out on the LTQ-XL equipped with an ESI-LC system Vanquish (ThermoFisher Scientific, Courtaboeuf, France). Peptides were separated using an Acclaim RSLC C_18_ column (2.2 μm; 2.1 × 150 mm; Thermofisher, France). The mobile phase was a gradient prepared from 0.1% aqueous formic acid (solvent A) and 0.1% formic acid in acetonitrile (solvent B). The peptides were eluted using a linear gradient from 0 to 50% of solvent B during 45 min, then from 50 to 100% during 10 min, and finally held for 5 min at a 250 μL min^-1^ flow rate. The electrospray ionization mass spectrometry detection was performed in positive mode with the following optimized parameters: the capillary temperature was set at 300°C, the spray voltage was +4.5 kV, and the sheath gas and auxiliary gas were set at 50 and 10 psi, respectively. The acquisition range was from 100 to 2000 *m/z*. The area value of each peak corresponding to a peptide was manually integrated using the peak ion extraction function in Xcalibur software (version 4.0, ThermoFisher Scientific, Courtaboeuf, France). The relative peak area indicates the contribution of each peptide to all the peptides identified in the venom, providing a measure of relative abundance.

For reduction/alkylation, 300 μg of crude venom was incubated with 10 μL of 100 mM ammonium bicarbonate buffer (pH 8) containing 10 mM dithiothreitol (DTT) for 30 min at 56°C. After reducing with DTT, the samples were alkylated by adding 10 μL of 50 mM iodoacetamide (IA) for 15 min at room temperature in the dark. Reduced/alkylated venoms were freeze-dried prior to shotgun proteomics. The venoms were resuspended in 100 μL 10% ACN and desalted using ZipTip μ-C_18_ Pipette Tips (Pierce Biotechnology, Rockford, IL, USA). Samples were analyzed using an Orbitrap Fusion equipped with an easy spray ion source and coupled to a nano-LC Proxeon 1200 (Thermo Scientific, Waltham, MA, USA). Peptides were loaded with an online preconcentration method and separated by chromatography using a Pepmap-RSLC C_18_ column (0.75 × 750 mm, 2 μm, 100 Å) from Thermo Scientific, equilibrated at 50°C and operating at a flow rate of 300 nL/min. Peptides were eluted by a gradient of solvent A (H_2_O, 0.1% FA) and solvent B (ACN/H_2_O 80/20, 0.1% FA), the column was first equilibrated 5 min with 95% of A, then B was raised to 28% in 105 min and to 40% in 15 min. Finally, the column was washed with 95% B during 20 min and re-equilibrated at 95% A during 10 min. The Advanced Peak Determination (APD) algorithm was used during the acquisition. Peptide masses were analyzed in the Orbitrap cell in full ion scan mode, at a resolution of 120,000, a mass range of *m/z* 350-1550 and an AGC target of 4.105. MS/MS were performed in the top speed 3 s mode. Peptides were selected for fragmentation by Higher-energy C-trap Dissociation (HCD) with a Normalized Collisional Energy of 27% and a dynamic exclusion of 60 s. Fragment masses were measured in an Ion trap in the rapid mode, with an AGC target of 1.104. Monocharged peptides and unassigned charge states were excluded from the MS/MS acquisition. The maximum ion accumulation times were set to 100 ms for MS and 35 ms for MS/MS acquisitions respectively. Using PEAKS X+ Studio (Bioinformatics Solutions Inc., Waterloo, ON, Canada) in no-enzyme mode, MS/MS spectra were searched against the translated venom-apparatus transcriptomes. Precursor and fragment mass tolerances were set to respectively 7 ppm and 0.5 Da. The following post-translational modifications were included as variable: Acetyl (Protein N-term), Oxidation (M), Phosphorylation (STY), Deamidation (NQ), Amidation (C-term), HexNAcylation (ST), Pyro-glu (EQ). The following post-translational modifications were included as fixed: Carbamidomethyl (C). Spectra were filtered using a 1% FDR.

### Venom composition analysis

To characterize the venom composition of each species we employed a transcriptomic approach with mass spectrometry to validate the presence of peptides in the venom. From the transcriptomes generated, the annotations were manually curated focusing on transcripts coding for toxins, with consideration of gene expression levels in venom glands and in the body. Additionally, we selected and examined additional transcripts based on precursor similarities to known toxin peptides. A total of 465 transcripts encoding putative toxins were retained for subsequent analysis (***SI Appendix*, Figure S8** and **Dataset S1**). Crude venom from all species was obtained by dissection of venom reservoirs. Venoms were then reduced/alkylated and analyzed through shotgun LC-MS/MS proteomics. The PEAKS software was used to analyze the mass spectrometry fragmentation spectra, with the transcriptome of each species implemented as a database for peptide sequence assignment. Positive matches of proteomics data with transcripts allowed us to confirm 305 peptide sequences. Total ion chromatograms were also generated with crude venoms and the LC-MS profile annotated **(*SI Appendix*, Figure S9-10)**.

Transcripts were classified into families based on the similarity of the amino acid sequences of the mature regions. For each family, multiple alignments of full-length precursors were generated using the Muscle program in MEGA-X version 10.1.8 (67) and edited using Jalview version 2.11.2.7 (68). We classified the transcripts into 62 peptide families and 3 enzyme toxins (i.e. phospholipase A_1_, phospholipase A_2_, and venom allergen 3) **(*SI Appendix*, Figure S11-18**). Peptide transcripts were further clustered into five gene clades (i.e. cysteine-rich poneritoxin, ponericin-like, pseudomyrmeciitoxin (PSDTX), ICK-PONTX, dimeric myrmicitoxin (MYRTX)) according to the predictive signal sequence (***SI Appendix*, Figure S19**). A total of 14 toxin families (i.e. families 23, 26, 29, 33, 34, 35, 37, 49, 52, 54, 56, and 59) were not included in the venom composition analysis for clustering because no transcript sequences could be confirmed by mass spectrometry, yet they were retained in the sequence alignments. The full list of transcripts expressed in venom glands with a TMM greater than 100, the identified toxin precursor sequence, the predicted mature part, the family assignment, and PEAKS results can be found in **Dataset S1**. The venom composition of *Pa. clavata*, previously published by Aili *et al* (46) has been included in our analysis. For venom clustering, a Bray-Curtis distance matrix based on the relative expression of the toxin family was generated using the veggdist () function from the R package “vegan” (69), followed by hierarchical clustering analysis (HCA) using the hclust () function with the full method from the R package “stats” (70). The dendlist () function from the R package “dendextend” (71) was used to plot and align the species phylogeny tree with the venom composition HCA cladogram. The final Figure 2 was edited in GraphPad Prism v10.0.3. To define gene clades, we used an approach based on similarity of signal parts. Signal parts were predicted using SignalP - 6.0 server (72) and then aligned using the Muscle program in MEGA-X (67). A pairwise distance matrix between sequences was extracted from the multiple alignments and used for HCA clustering using the hclust () function with the ward method from the R package “stats”.

### Morphological traits

Six morphological traits were measured on up to 13 randomly selected workers per species. Measurements were made using an ocular micrometer accurate to 0.01 mm mounted on a Leica M80 or Leica S9E stereomicroscope (Leica Microsystems, Heerbrugg, Switzerland). The traits considered were Weber’s length, head length, mandible length, sting length, venom reservoir length, and venom reservoir width. We estimated the venom reservoir volume using the standard ellipsoid formula; π/6 (venom reservoir length × venom reservoir width^2^). For analysis we used size/volume-corrected ratios calculated as follows: mandible proportion (mandible length / head length), sting proportion (sting length / weber’s length), venom reservoir proportion (venom reservoir volume / weber’s length^3^).

### Neuronal cells assays

F11 (mouse neuroblastoma × DRG neuron hybrid) were maintained on Ham’s F12 media supplemented with 10% FBS, 100 μM hypoxanthine, 0.4 μM aminopterin, and 16 μM thymidine (Hybri-Max, Sigma Aldrich). 384-well imaging plates (Corning, Lowell, MA, USA) were seeded 24 h prior to calcium imaging, resulting in ∼90% confluence at the time of imaging. Cells were loaded for 30 min at 37°C with Calcium 4 assay component A in physiological salt solution (PSS; 140 mM NaCl, 11.5 mM D-glucose, 5.9 mM KCl, 1.4 mM MgCl_2_, 1.2 mM NaH_2_PO_4_, 5 mM NaHCO_3_, 1.8 mM CaCl_2_, 10 mM HEPES) according to the manufacturer’s instructions (Molecular Devices, Sunnyvale, CA). Ca^2+^ responses were measured using a FLIPRPenta fluorescent plate reader equipped with a CCD camera (Ex: 470 to 490 nm, Em: 515 to 575 nM) (Molecular Devices, Sunnyvale, CA). Signals were read every second for 10 s before, and 300 s after, the addition of venoms (in PSS supplemented with 0.1% BSA).

### Insect activity assays

Blowfly larvae (*Lucilia caesar*) were purchased from a fisheries shop (Euroloisir81, Lescure-d’Albigeois, France) and kept at 25°C until hatching. Flies 1-4 days after hatching were used for injection assays. Blowfly assays were done through lateral intrathoracic injection of 1 μL of venom dissolved in ultra-pure water at various concentrations using a fixed 25-gauge needle attached to an Arnold hand microapplicator (Burkard Manufacturing Co., Ltd., Rickmansworth, UK) with a 1.0 mL Hamilton Syringe (1000 Series Gastight, Hamilton Company, Reno, NV, USA).Then, the blowfly was placed in an individual 2 mL tube containing 15 μL of 5% glucose solution. Paralysis was monitored at 1 h and 24 h post-injection, while lethality was monitored at 24 h. Flies that did not display any signs of movement dysfunction were considered unaffected, otherwise they were recorded as paralyzed. Flies were deemed dead if they did not respond at all to tweezer mechanical stimulation as observed under a dissecting microscope. Ten flies were used for each toxicity experiment and for the corresponding control (ultrapure water solution). Each dose was repeated three times.

### Cytotoxicity and membrane potential assays

*Drosophila* S2 cells (Thermofisher, USA) were maintained and prepared for cytotoxicity and membrane potential variation assays as previously detailed (34, 73). Lyophilized crude venoms were solubilized in ultra-pure water and diluted in culture medium before being exposed to cells at various final concentrations (from 1 ng/mL to 100 μg/mL) for 24 h at 25°C for cytotoxic assays or at 100 μg/ml for 30 min at 25°C for membrane potential monitoring. Cytotoxic assays were performed using lysis buffer and culture medium as positive and negative controls or blanks, respectively. The assays and calculations of LC_50_ were performed as previously described (73). Membrane potential changes were measured using a buffer containing: 115 mM NaCl, 5 mM KCl, 2 mM CaCl_2_, 1 mM MgCl_2_, 48 mM sucrose, and 10 mM HEPES. The assay and analysis were performed as previously described (34).

### Phylogenetic analysis

To generate a phylogeny of the studied species we searched for conserved genes across their transcriptome assemblies, or genome for *Pa. clavata*, using BUSCO v5.1.2 (59). Using the hymenoptera_odb10 database, we identified 566 genes present in at least 90 % of the species. We aligned the protein sequences from each gene using mafft and concatenated them into a supermatrix (**Dataset S2**). Then, we used IQTREE2 v2.1.2 (74) to reconstruct the maximum likelihood tree by using the concatenated matrix and the selection of the best substitution model with ModelFinder and 1000 ultrafast bootstrap replicates.

Ancestral states reconstruction for venom activities and capacities, were estimated by maximum likelihood using the contmap () function of the Phytools R package with default settings. To test for statistical support for correlations between venom bioactivities and morphological traits, and the influences of ecological traits, we used the phylogenetic Generalized Least Squares (PGLS) approach using the PGLS () function of the “caper” package in RStudio, with the formula set as ([functional traits, e.g. vertebrate_pain] ∼ grouping [ecological traits, e.g. defensive_use (yes or no)] and lambda set to “Maximum Likelihood”. For the PGLS regressions, we treated log-transformed continuous traits.

## Acknowledgments

We thank Wolfgang Wuster and Nicholas Casewell for valuable input into the experimental design. We thank Philippe Gaucher for providing a colony of *Pseudomyrmex viduus*. We thank Federica Catonaro, Elena di Barbora, and Emanuela Aleo for assistance with transcriptome library construction and sequencing. This research was funded by Investissement d’Avenir grant of the Agence Nationale de la Recherche (CEBA: ANR-10-LABX-25-01) and by the PO-FEDER 2014–2020, Région Guyane (FORMIC, GY0013708). Ant samples were collected under the authorizations of the French Ministry of Ecological and Solidarity Transition, in accordance with Article 17, paragraph 2, of the Nagoya Protocol on Access and Benefit-sharing (Reference number of the permit: TREL1916196S/214).

## Data availability

The raw sequencing reads used in this manuscript are available from the National Center for Biotechnology Information (NCBI) under the project code PRJNA1061791. The mass spectrometry proteomics data have been deposited to the ProteomeXchange Consortium via the PRIDE partner repository with the dataset identifier PXD050348.

## Author contributions

A.T., J.O., and N.T. designed research; A.T., S.D.R., H.L., S.A., N.J.T., V.T., F.P., and A.B. performed research; J.O., N.T., E.B., M.T., I.V., and C.S.M. contributed resources; A.T., S.D.R., and H.L. analyzed data; A.T. wrote the manuscript; all authors have read and agreed to the published version of the manuscript.

## References

1. N. R. Casewell, W. Wüster, F. J. Vonk, R. A. Harrison, B. G. Fry, Complex cocktails: the evolutionary novelty of venoms. Trends Ecol. Evol. 28, 219–229 (2013).

2. Z. Nisani, W. K. Hayes, Defensive stinging by Parabuthus transvaalicus scorpions: risk assessment and venom metering. Anim. Behav. 81, 627–633 (2011).

3. S. Dutertre, et al., Evolution of separate predation- and defence-evoked venoms in carnivorous cone snails. Nat. Commun. 5, 3521 (2014).

4. E. R. J. Evans, T. D. Northfield, N. L. Daly, D. T. Wilson, Venom costs and optimization in scorpions. Frontiers in Ecology and Evolution 7 (2019).

5. S. D. Robinson, et al., A comprehensive portrait of the venom of the giant red bull ant, Myrmecia gulosa, reveals a hyperdiverse hymenopteran toxin gene family. Sci Adv 4, eaau4640 (2018).

6. V. Schendel, L. D. Rash, R. A. Jenner, E. A. B. Undheim, The diversity of venom: The importance of behavior and venom system morphology in understanding its ecology and evolution. Toxins 11 (2019).

7. B. D. Blanchard, C. S. Moreau, Defensive traits exhibit an evolutionary trade-off and drive diversification in ants. Evolution 71, 315–328 (2017).

8. A. Touchard, et al., The biochemical toxin arsenal from ant venoms. Toxins 8 (2016).

9. E. O. Wilson, B. Hölldobler, The rise of the ants: a phylogenetic and ecological explanation. Proc. Natl. Acad. Sci. U. S. A. 102, 7411–7414 (2005).

10. X. Cerdá, A. Dejean, Predation by ants on arthropods and other animals. National Academy of Sciences (US) (2011).

11. D. B. Booher, et al., Functional innovation promotes diversification of form in the evolution of an ultrafast trap-jaw mechanism in ants. PLoS Biol. 19, e3001031 (2021).

12. F. J. Larabee, A. V. Suarez, The evolution and functional morphology of trap-jaw ants (Hymenoptera: Formicidae). Myrmecol. News 20, 25–36 (2014).

13. J. Orivel, A. Dejean, Comparative effect of the venoms of ants of the genus Pachycondyla (Hymenoptera: Ponerinae). Toxicon 39, 195–201 (2001).

14. B. E. R. Rubin, S. Kautz, B. D. Wray, C. S. Moreau, Dietary specialization in mutualistic acacia-ants affects relative abundance but not identity of host-associated bacteria. Mol. Ecol. 28, 900–916 (2019).

15. C. Jelley, C. S. Moreau, Aggressive behavior across ant lineages: importance, quantification, and associations with trait evolution. Insectes Soc. (2023).

16. D. C. Cardoso, Í. C. C. Alves, M. P. Cristiano, J. Heinze, Death feigning in ants. Myrmecol. News 34 (2024).

17. D. A. Grasso, D. Giannetti, C. Castracani, F. A. Spotti, A. Mori, Rolling away: a novel context-dependent escape behaviour discovered in ants. Sci. Rep. 10, 3784 (2020).

18. V. Barassé, et al., Venomics survey of six myrmicine ants provides insights into the molecular and structural diversity of their peptide toxins. Insect Biochem. Mol. Biol. 151, 103876 (2022).

19. A. Touchard, et al., The complexity and structural diversity of ant venom peptidomes is revealed by mass spectrometry profiling. Rapid Commun. Mass Spectrom. 29, 385–396 (2015).

20. T. D. Kazandjian, et al., Convergent evolution of pain-inducing defensive venom components in spitting cobras. Science 371, 386–390 (2021).

21. A. E. Mill, Predation by the ponerine ant Pachycondyla commutata on termites of the genus Syntermes in Amazonian rain forest. J. Nat. Hist. 18, 405–410 (1984).

22. A. Dejean, I. Olmsted, Ecological studies on Aechmea bracteata (Swartz) (Bromeliaceae). J. Nat. Hist. 31, 1313–1334 (1997).

23. B. Schatz, J. Orivel, J.-P. Lachaud, G. Beugnon, A. Dejean, Sitemate recognition: the case of Anochetus traegordhi (Hymenoptera; Formicidae) preying on Nasutitermes (Isoptera: Termitidae). Sociobiology 34, 569–580 (1999).

24. A. Dejean, N. Labrière, A. Touchard, F. Petitclerc, O. Roux, Nesting habits shape feeding preferences and predatory behavior in an ant genus. Naturwissenschaften 101, 323–330 (2014).

25. T. Piek, et al., Poneratoxin, a novel peptide neurotoxin from the venom of the ant, Paraponera clavata. Comp. Biochem. Physiol. C 99, 487–495 (1991).

26. A. Camargo, PCAtest: testing the statistical significance of Principal Component Analysis in R. PeerJ 10, e12967 (2022).

27. L. J. Revell, phytools: an R package for phylogenetic comparative biology (and other things). Methods Ecol. Evol. 3, 217–223 (2012).

28. C. Schmidt, Molecular phylogenetics of ponerine ants (Hymenoptera: Formicidae: Ponerinae). Zootaxa 3647, 201–250 (2013).

29. N. F. Carlin, D. S. Gladstein, The “bouncer” defense of Odontomachus Ruginodis and other odontomachine ants (Hymenoptera: Formicidae). Psyche 96, 1–19 (1989).

30. A. Dejean, et al., The ecology and feeding habits of the arboreal trap-jawed ant Daceton armigerum. PLoS One 7, e37683 (2012).

31. E. Van Wilgenburg, M. A. Elgar, Colony characteristics influence the risk of nest predation of a polydomous ant by a monotreme. Biol. J. Linn. Soc. Lond. 92, 1–8 (2007).

32. A. De la Mora, G. Pérez-Lachaud, J.-P. Lachaud, Mandible strike: the lethal weapon of Odontomachus opaciventris against small prey. Behav. Processes 78, 64–75 (2008).

33. J. C. Gibson, F. J. Larabee, A. Touchard, J. Orivel, A. V. Suarez, Mandible strike kinematics of the trap-jaw ant genus Anochetus Mayr (Hymenoptera: Formicidae). J. Zool. 306, 119–128 (2018).

34. V. Barassé, et al., Discovery of an insect neuroactive helix ring peptide from ant venom. Toxins 15 (2023).

35. A. Touchard, et al., Isolation and characterization of a structurally unique β-hairpin venom peptide from the predatory ant Anochetus emarginatus. Biochim. Biophys. Acta 1860, 2553–2562 (2016).

36. J. O. Schmidt, Pain and Lethality Induced by Insect Stings: An Exploratory and Correlational Study. Toxins 11 (2019).

37. K. Kazuma, K. Masuko, K. Konno, H. Inagaki, Combined venom gland transcriptomic and venom peptidomic analysis of the predatory ant Odontomachus monticola. Toxins 9, 323 (2017).

38. J. Orivel, et al., Ponericins, new antibacterial and insecticidal peptides from the venom of the ant Pachycondyla goeldii. J. Biol. Chem. 276, 17823–17829 (2001).

39. S. R. Johnson, J. A. Copello, M. S. Evans, A. V. Suarez, A biochemical characterization of the major peptides from the Venom of the giant Neotropical hunting ant Dinoponera australis. Toxicon 55, 702–710 (2010).

40. S. Lv, et al., Highly selective performance of rationally designed antimicrobial peptides based on ponericin-W1. Biomater Sci 10, 4848–4865 (2022).

41. A. S. Senetra, M. R. Necelis, G. A. Caputo, Investigation of the structure-activity relationship in ponericin L1 from Neoponera goeldii. Pept. Sci. 112 (2020).

42. S. A. Nixon, et al., Multipurpose peptides: The venoms of Amazonian stinging ants contain anthelmintic ponericins with diverse predatory and defensive activities. Biochem. Pharmacol. 192, 114693 (2021).

43. T. Jensen, et al., Venom chemistry underlying the painful stings of velvet ants (Hymenoptera: Mutillidae). Cell. Mol. Life Sci. 78, 5163–5177 (2021).

44. A. A. Walker, et al., Production, composition, and mode of action of the painful defensive venom produced by a limacodid caterpillar, Doratifera vulnerans. Proc. Natl. Acad. Sci. U. S. A. 118 (2021).

45. Z. Dekan, et al., Δ-Myrtoxin-Mp1a is a helical heterodimer from the venom of the jack jumper ant that has antimicrobial, membrane-disrupting, and nociceptive activities. Angew. Chem. Weinheim Bergstr. Ger. 129, 8615–8619 (2017).

46. S. R. Aili, et al., An integrated proteomic and transcriptomic analysis reveals the venom complexity of the bullet ant Paraponera clavata. Toxins 12 (2020).

47. S. D. Robinson, et al., Ant venoms contain vertebrate-selective pain-causing sodium channel toxins. Nat. Commun. 14, 2977 (2023).

48. S. D. Robinson, et al., Peptide toxins that target vertebrate voltage-gated sodium channels underly the painful stings of harvester ants. J. Biol. Chem. 300, 105577 (2024).

49. J. Pan, W. F. Hink, Isolation and characterization of myrmexins, six isoforms of venom proteins with anti-inflammatory activity from the tropical ant, Pseudomyrmex triplarinus. Toxicon 38, 1403–1413 (2000).

50. A. Touchard, et al., Heterodimeric insecticidal peptide provides new insights into the molecular and functional diversity of ant venoms. ACS Pharmacol Transl Sci 3, 1211–1224 (2020).

51. P. A. Koenig, C. S. Moreau, Testing optimal defence theory in a social insect: Increased risk is correlated with increased venom investment. Ecol. Entomol. (2023) 10.1111/een.13295.

52. M. Hölzer, M. Marz, De novo transcriptome assembly: A comprehensive cross-species comparison of short-read RNA-Seq assemblers. Gigascience 8 (2019).

53. Y. Xie, et al., SOAPdenovo-Trans: de novo transcriptome assembly with short RNA-Seq reads. Bioinformatics 30, 1660–1666 (2014).

54. E. Bushmanova, D. Antipov, A. Lapidus, A. D. Prjibelski, rnaSPAdes: a de novo transcriptome assembler and its application to RNA-Seq data. Gigascience 8 (2019).

55. M. G. Grabherr, et al., Full-length transcriptome assembly from RNA-Seq data without a reference genome. Nat. Biotechnol. 29, 644–652 (2011).

56. M. Martin, Cutadapt removes adapter sequences from high-throughput sequencing reads. EMBnet.journal 17, 10–12 (2011).

57. B. E. R. Rubin, C. S. Moreau, Comparative genomics reveals convergent rates of evolution in ant– plant mutualisms. Nat. Commun. 7, 1–11 (2016).

58. R. Smith-Unna, C. Boursnell, R. Patro, J. M. Hibberd, S. Kelly, TransRate: reference-free quality assessment of de novo transcriptome assemblies. Genome Res. 26, 1134–1144 (2016).

59. F. A. Simão, R. M. Waterhouse, P. Ioannidis, E. V. Kriventseva, E. M. Zdobnov, BUSCO: Assessing genome assembly and annotation completeness with single-copy orthologs. Bioinformatics 31, 3210–3212 (2015).

60. E. Bushmanova, D. Antipov, A. Lapidus, V. Suvorov, A. D. Prjibelski, rnaQUAST: a quality assessment tool for de novo transcriptome assemblies. Bioinformatics 32, 2210–2212 (2016).

61. B. J. Haas, et al., De novo transcript sequence reconstruction from RNA-seq using the Trinity platform for reference generation and analysis. Nat. Protoc. 8, 1494–1512 (2013).

62. H. Nielsen, Predicting Secretory Proteins with SignalP. Methods Mol. Biol. 1611, 59–73 (2017).

63. A. Krogh, B. Larsson, G. von Heijne, E. L. Sonnhammer, Predicting transmembrane protein topology with a hidden Markov model: application to complete genomes. J. Mol. Biol. 305, 567–580 (2001).

64. B. Langmead, S. L. Salzberg, Fast gapped-read alignment with Bowtie 2. Nat. Methods 9, 357–359 (2012).

65. B. Li, C. N. Dewey, RSEM: accurate transcript quantification from RNA-Seq data with or without a reference genome. BMC Bioinformatics 12, 323 (2011).

66. D. M. Bryant, et al., A Tissue-Mapped Axolotl De Novo Transcriptome Enables Identification of Limb Regeneration Factors. Cell Rep. 18, 762–776 (2017).

67. S. Kumar, G. Stecher, M. Li, C. Knyaz, K. Tamura, MEGA X: Molecular Evolutionary Genetics Analysis across Computing Platforms. Mol. Biol. Evol. 35, 1547–1549 (2018).

68. A. M. Waterhouse, J. B. Procter, D. M. A. Martin, M. Clamp, G. J. Barton, Jalview Version 2—a multiple sequence alignment editor and analysis workbench. Bioinformatics 25, 1189–1191 (2009).

69. P. Dixon, VEGAN, a package of R functions for community ecology. J. Veg. Sci. 14, 927–930 (2003).

70. R. R Core Team, Others, R: A language and environment for statistical computing (2013).

71. T. Galili, dendextend: an R package for visualizing, adjusting and comparing trees of hierarchical clustering. Bioinformatics 31, 3718–3720 (2015).

72. F. Teufel, et al., SignalP 6.0 predicts all five types of signal peptides using protein language models. Nat. Biotechnol. 40, 1023–1025 (2022).

73. S. Ascoët, et al., The mechanism underlying toxicity of a venom peptide against insects reveals how ants are master at disrupting membranes. iScience 26, 106157 (2023).

74. B. Q. Minh, et al., IQ-TREE : New Models and Efficient Methods for Phylogenetic Inference in the Genomic Era. Mol. Biol. Evol. 37, 2461 (2020).

