## Supplementary material for "Adaptive trade-offs between vertebrate defense and insect predation drive ant venom evolution": SI Appendix

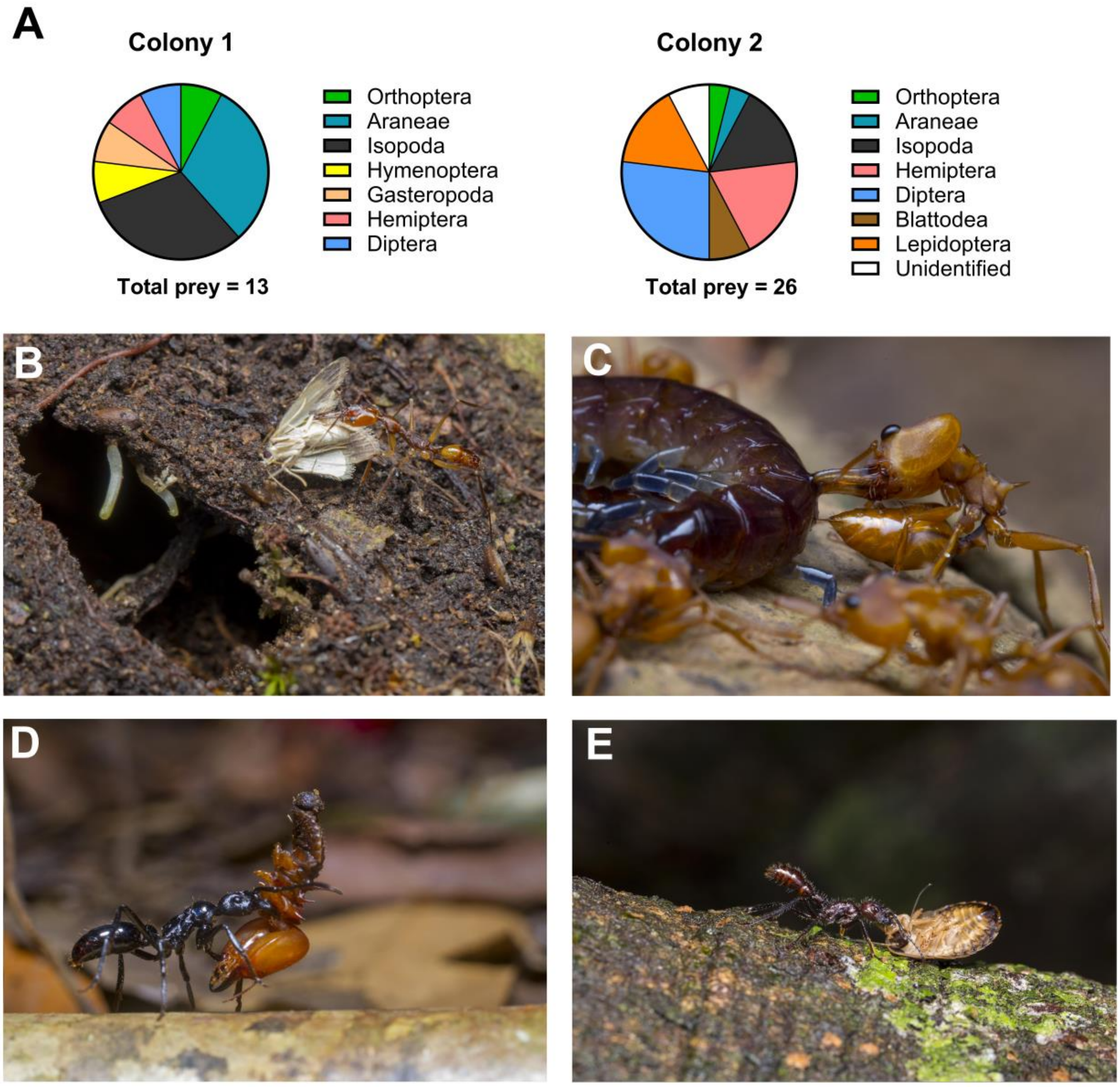


**Figure S1. Field observations of defensive and predatory behaviors of stinging ants. A)** We conducted *in situ* night-time observations of two colonies of *A. emarginatus* and collected all workers returning to the nests with prey in their mandibles. Morphological examination of the prey in the laboratory enabled us to identify 37 prey items at the order level, which included a diverse assortment of small arthropods (e.g. woodlice, spiders, cicadas, flies, moths). Pie charts showing the identified prey items of two colonies of *A. emarginatus*. **B)** *Anochetus emarginatus* bringing an immobilized moth prey into the nest. We observed that *A. emarginatus* systematically used its sting on termites following mandibular seizure. **C)** *Daceton armigerum* worker stinging a centipede on which it is preying. *Daceton armigerum* does not use its sting for defense but stings its prey, especially when it is large and difficult to handle. **D)** Immobilized *Syntermes* sp. termite being transported by *Neoponera commutata*. **E)** *Paraponera clavata* worker bringing a cockroach prey to the nest. Although there were no direct observations of predatory stings by *N. commutata* or *Pa. clavata*, all prey brought back to the colony by workers were immobile suggestive of a paralytic role of the venom.

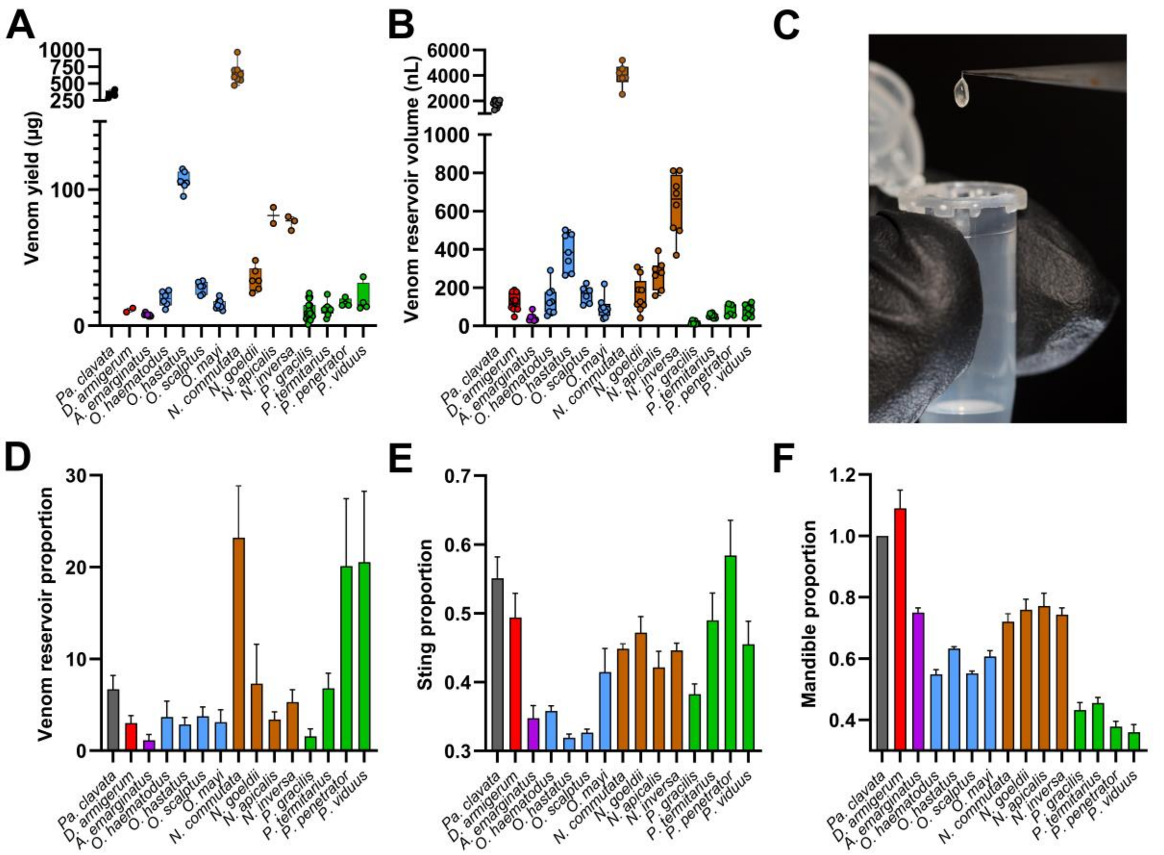


**Figure S2. Venom yield and morphological measurements. A)** Distribution of venom yield values per worker for each species obtained after dissection of venom reservoirs and lyophilization of filtered venom samples. **B)** Venom reservoir volume estimated by applying the formula for calculating the volume of an ellipsoid with the measured length and width of the venom reservoirs. **C)** Venom reservoir of *Neoponera commutata* obtained after dissection. Proportion of **D)** venom reservoir volume; **E)** sting length; and **F)** mandible. Data are presented as mean ± SEM (n = 4-13) (see values in **Table S1**).

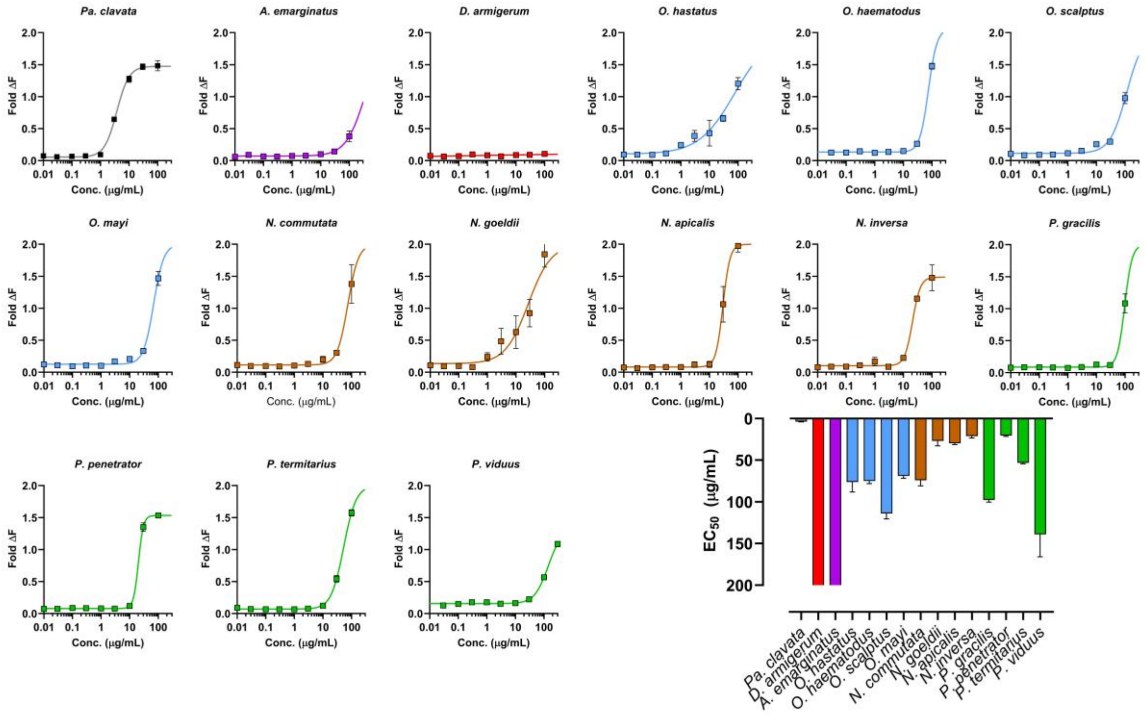


**Figure S3. Nocifensive activity of crude venoms.** Concentration-response curves of intracellular $Ca^{2+}$ concentration ($[Ca^{2+}]_i$) in the F11 (mouse neuroblastoma × rat dorsal root ganglion (DRG) neuron hybrid) cell line. Bar plot shows 50% maximal effective concentration ($EC_{50}$) for each species (see values in **Table S2**).

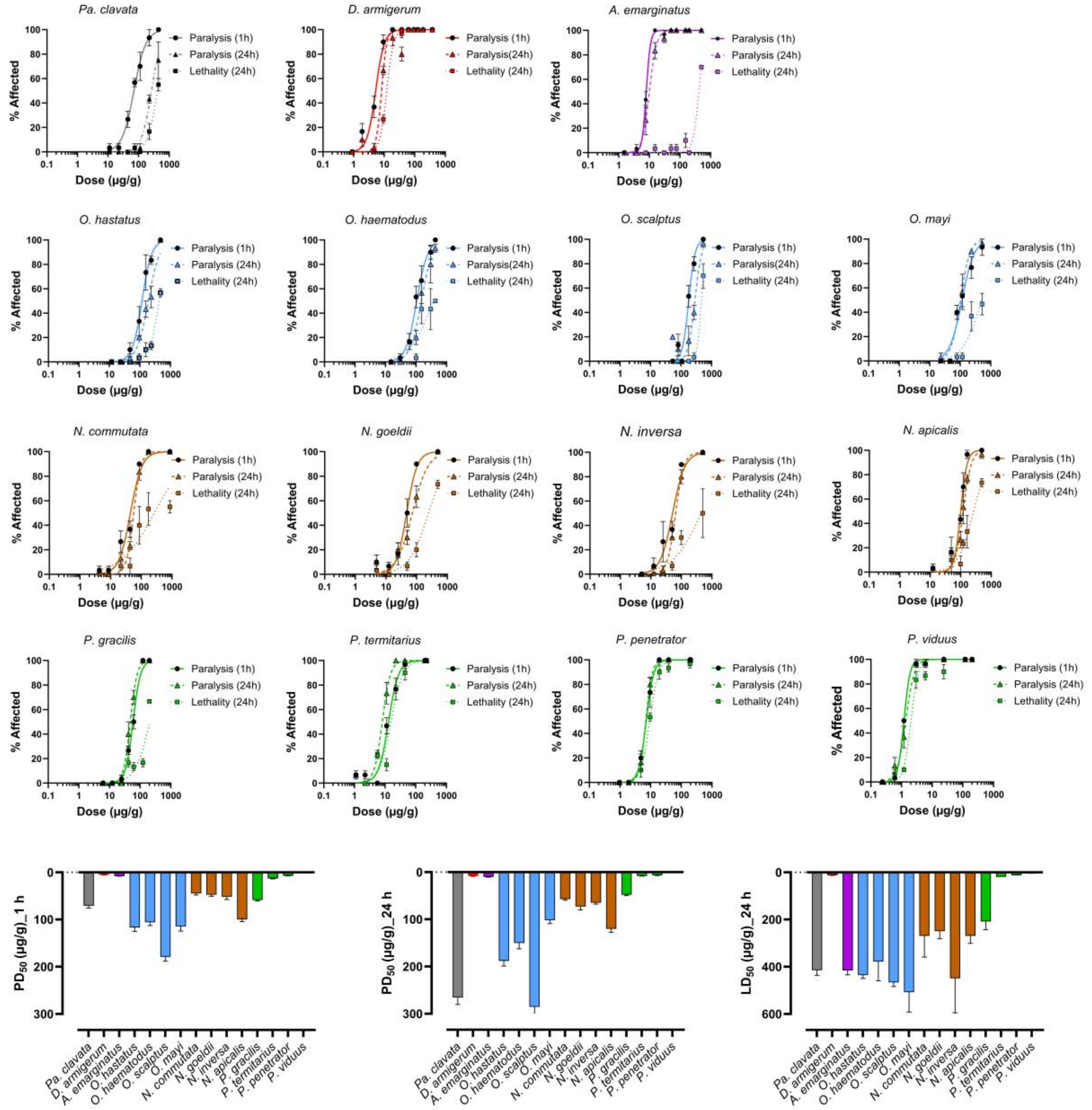


**Figure S4. Paralytic activity and lethality of crude venoms on the blowfly *Lucilia caesar*.** Dose-response curves for *L. caesar* blowflies (blue lines) injected with crude venoms, at 1 h and 24 h after intrathoracic injection. Values are the percentage of affected flies (paralyzed or dead) presented as mean ± SEM (n = 3 independent experiments) fitted with a non-linear regression with variable slope. Bar graphs show 50% paralytic dose ($PD_{50}$) and lethal dose ($DL_{50}$) values (see values in **Table S3**).

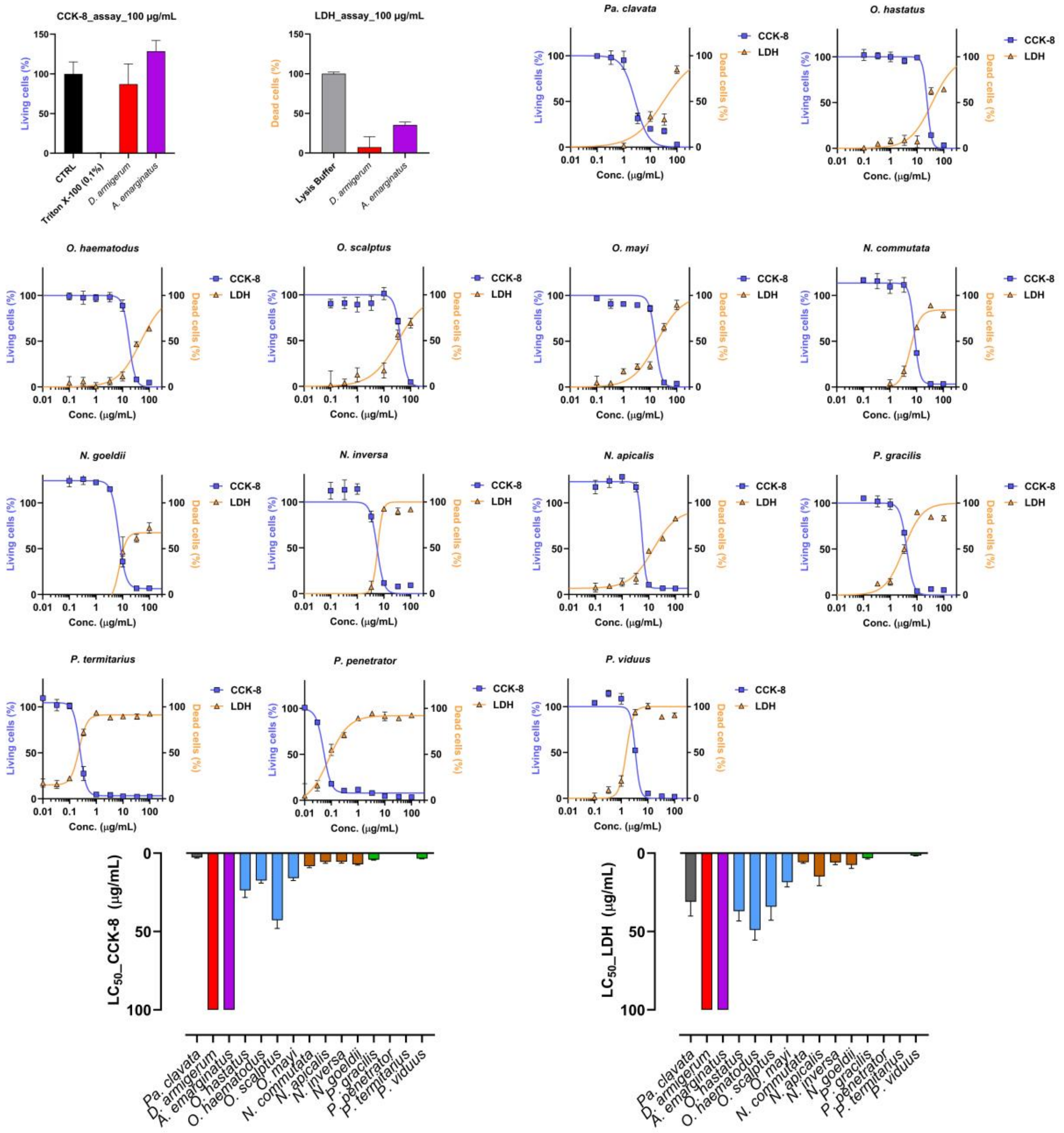


**Figure S5. Cytotoxic activity of crude venoms on *Drosophila* S2 cells obtained from CCK-8 and LDH assays.** No cytotoxic activity for *A. emarginatus* and *D. armigerum* crude venoms against S2 cells at 100 µg/mL. Concentration-response curves of *Neoponera* spp. *Odontomachus* spp., *Pseudomyrmex* spp., and *Pa. clavata* crude venoms against S2 cells. Values are means ± SEM (n = 1-3). The $LC_{50}$ of concentration-response curves were determined with a non-linear regression with variable slope (see values in **Table S4**).

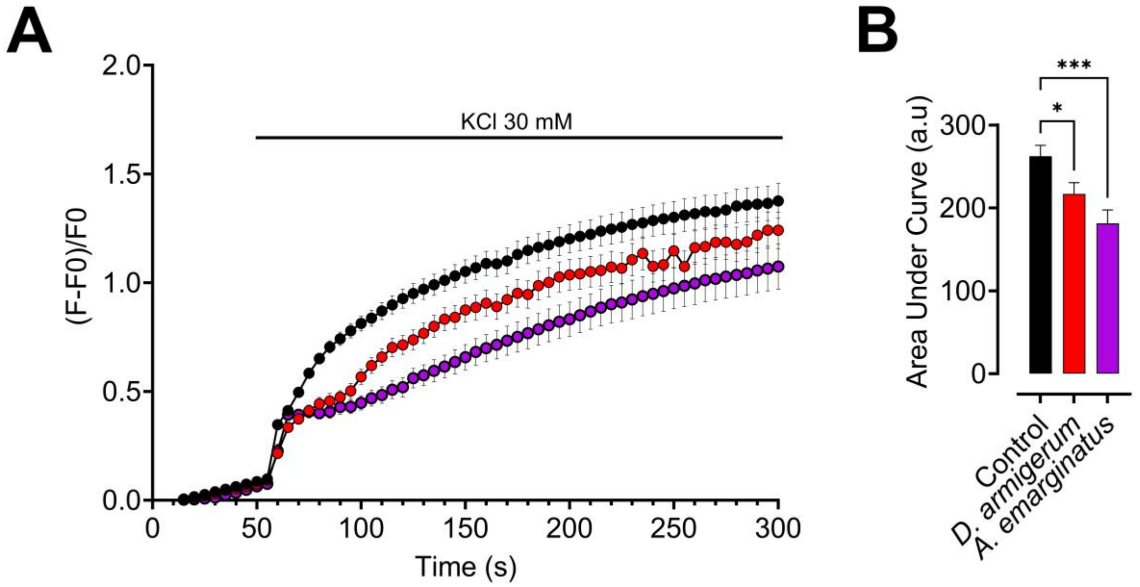


**Figure S6. Effect of non-cytotoxic crude venoms on *Drosophila* S2 cell membrane potential.** Cell membrane potential was recorded by DiBAC$_4$(3) after incubation with water (control) or 100 µg/ml crude venom. Membrane depolarization, indicated by an increase in relative fluorescence, was induced by application of high KCl concentration (30 mM). **A)** Time course of DiBAC$_4$(3) fluorescence in basal condition and after depolarization. **B)** Bar graph of the corresponding area under the curve (AUC). AUC were calculated after KCl application until the end of the experiment. Values are presented as mean SEM of N = 3 (with n = 142 for control condition, n = 90 cells for *D. armigerum* venom and n = 60 cells for *A. emarginatus* venom); *, $p < 0.05$ and ***, $p < 0.001$ compared to control condition (Welch's t-test).

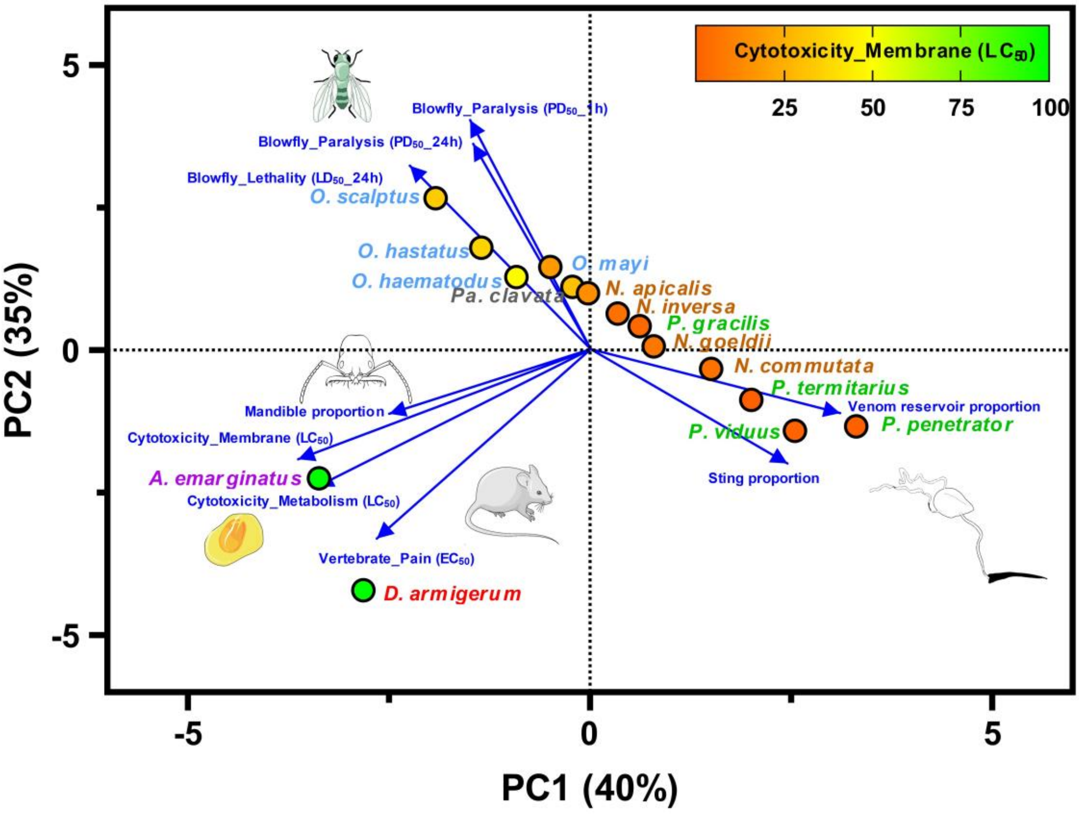


**Figure S7.** Principal component analysis of 15 ant species defined by venom bioactivities and morphological features.

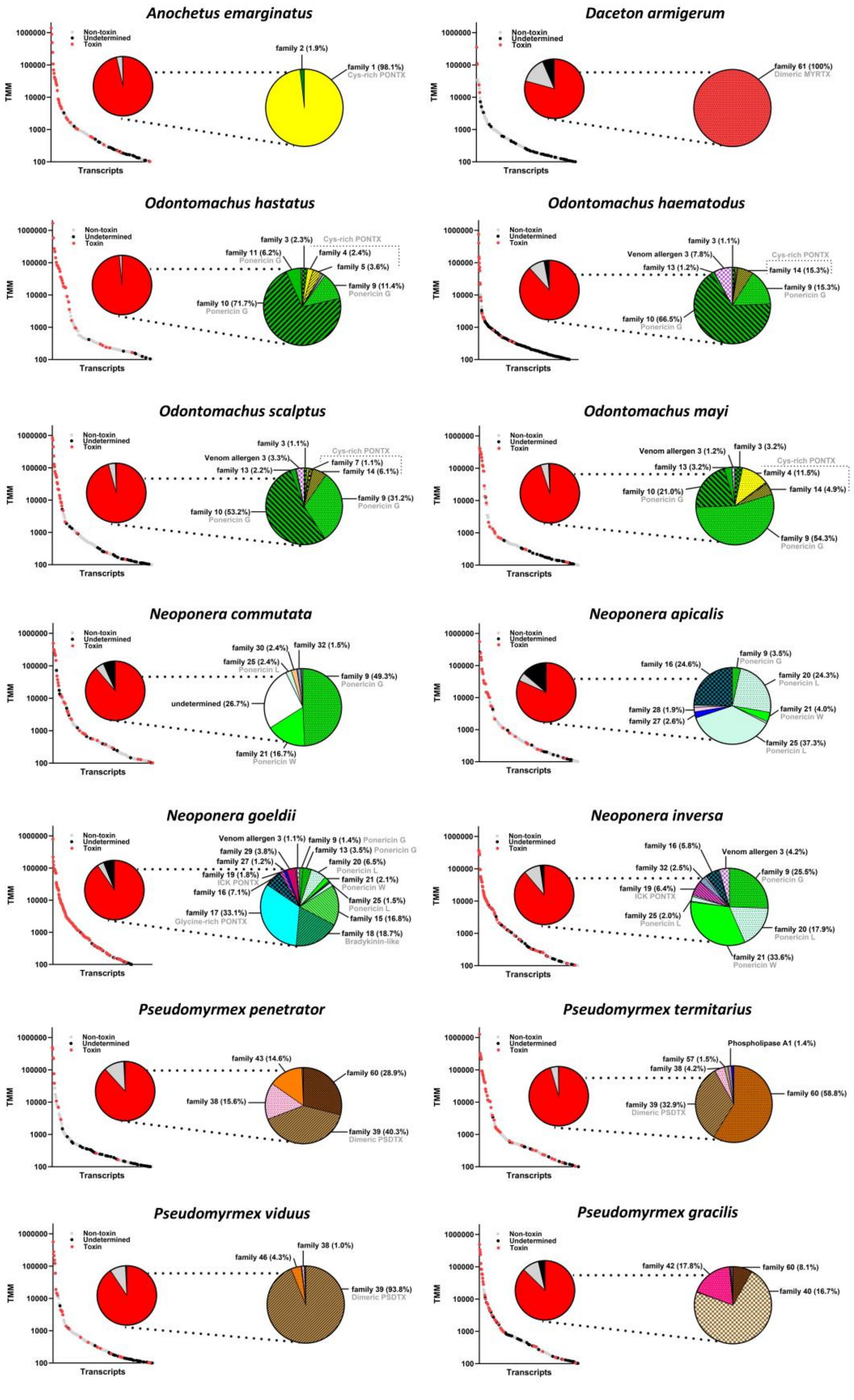


**Figure S8. Expression of transcripts encoding venom toxins (TMM values and relative expression in transcriptome).** The left of each panel shows the expression and relative proportion of transcripts expressed ≥ 100 TMM in the venom glands (with transcripts identified as toxin in red). The pie chart on the right of each panel shows the proportion of toxin families (only families with more than 1% relative expression are labeled). All peptide transcripts were classified into the toxin family based on sequence similarity.

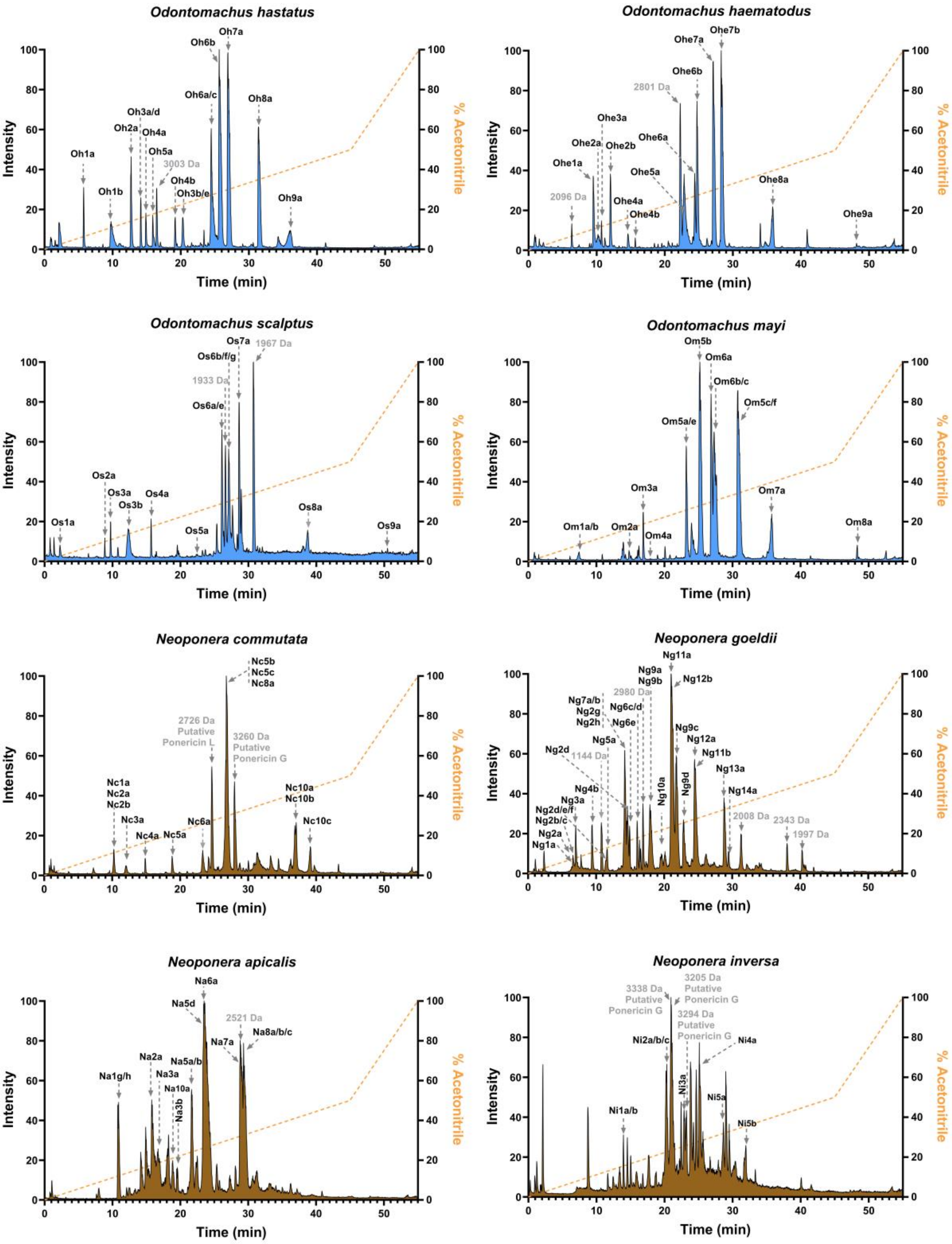


**Figure S9. Total ion chromatogram (TIC) of *Odontomachus* spp. and *Neoponera* spp. crude venoms.** HPLC peaks were labeled with the toxin ID when the measured masses in proteomic data matched the theoretical masses from the transcriptomic data.

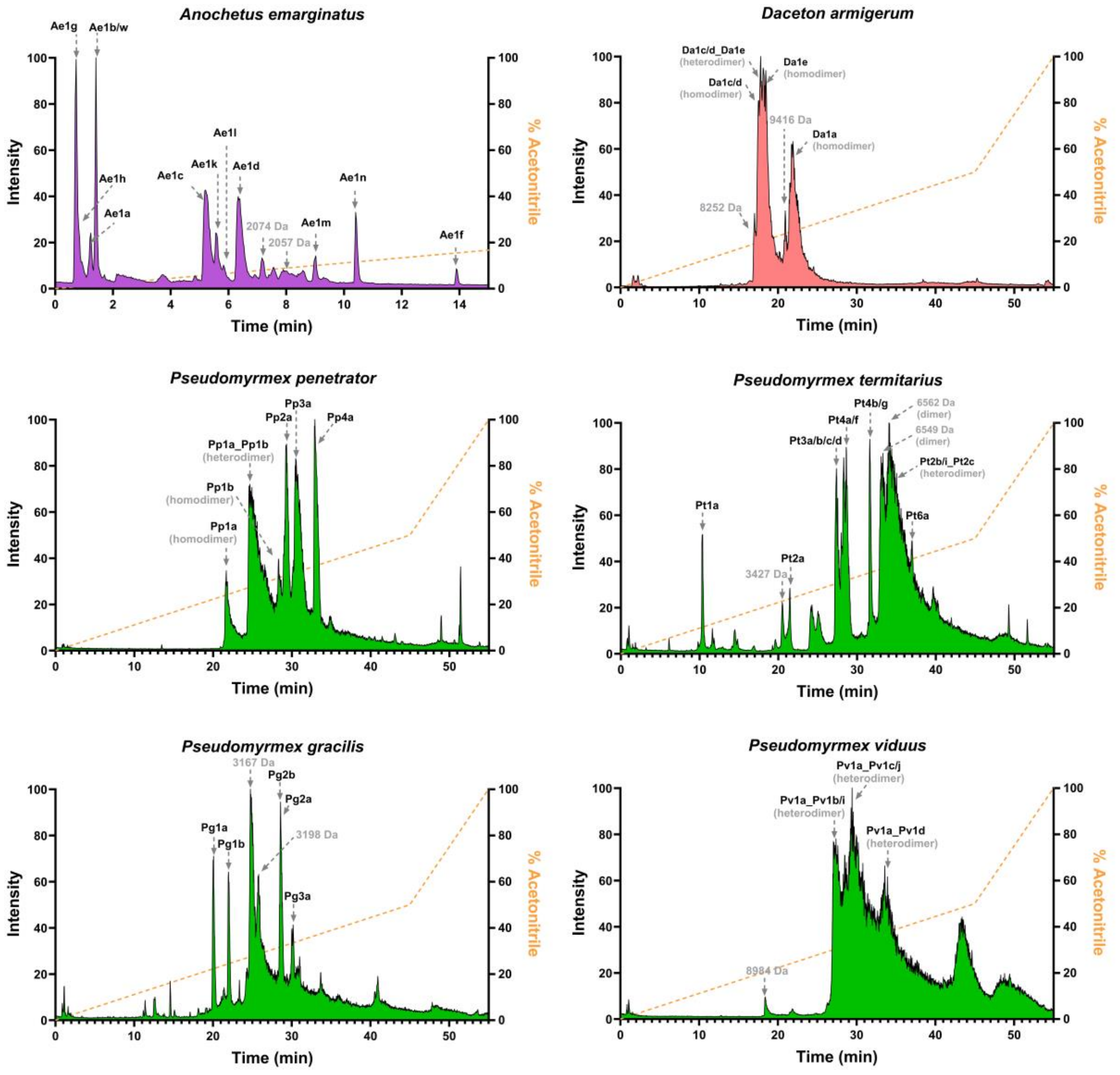


**Figure S10. Total ion chromatogram (TIC) of *Anochetus emarginatus, Daceton armigerum* and *Pseudomyrmex* spp. crude venoms.** HPLC peaks were labeled with the toxin ID when the measured masses in proteomic data matched the theoretical masses from the transcriptomic data.

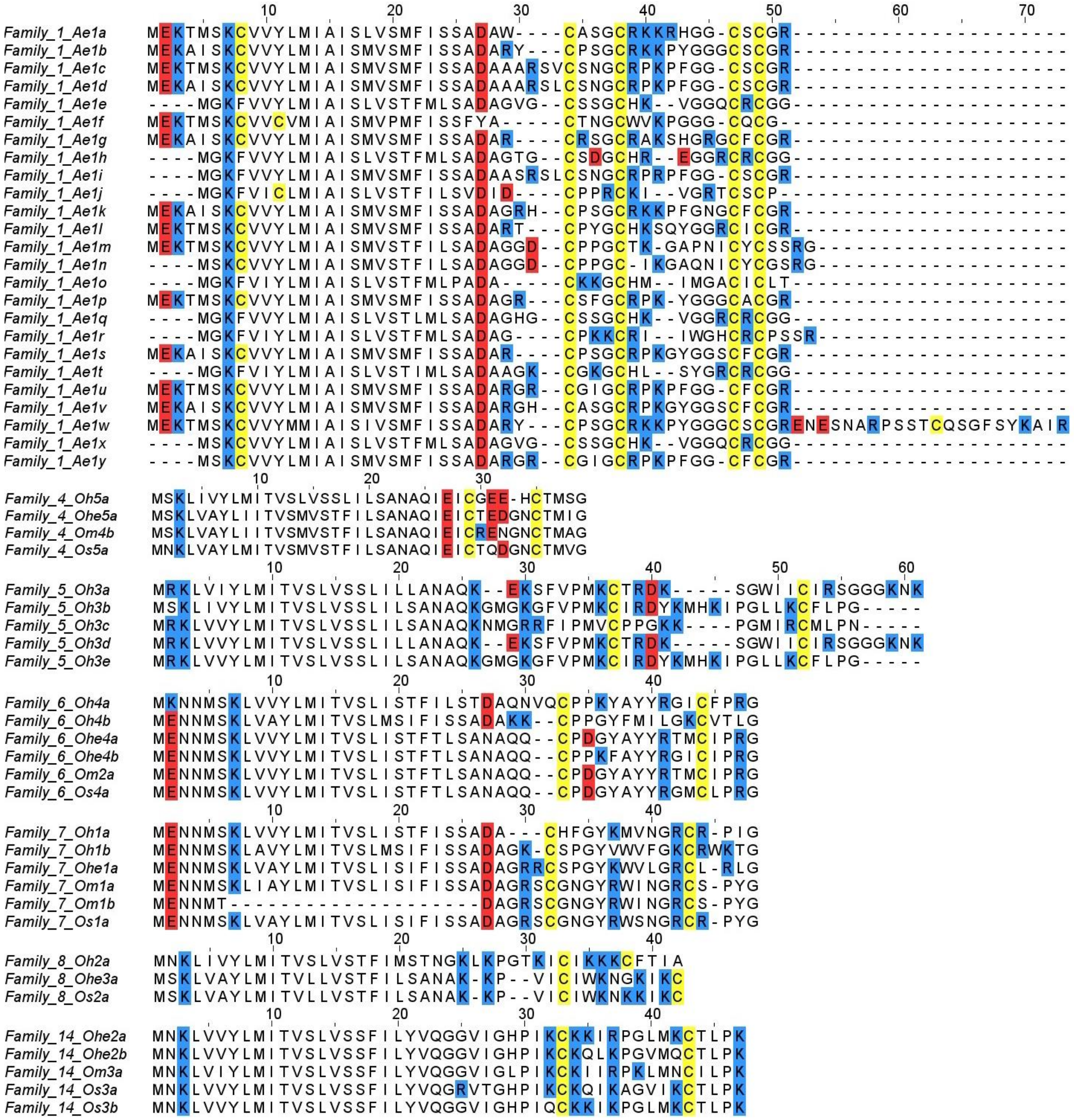

**Figure S11. Alignment of the amino acid sequences of toxin families belonging to the clade of Cysteine-rich Poneritoxins.** Multiple alignments were generated using the Muscle program in the MEGA-X version 10.1.8 and edited using Jalview version 2.11.2.7. Positively charged residues (lysine and arginine) are highlighted in blue, negatively charged residues (glutamic acid and aspartic acid) are highlighted in red, and cysteine residues are highlighted in yellow.

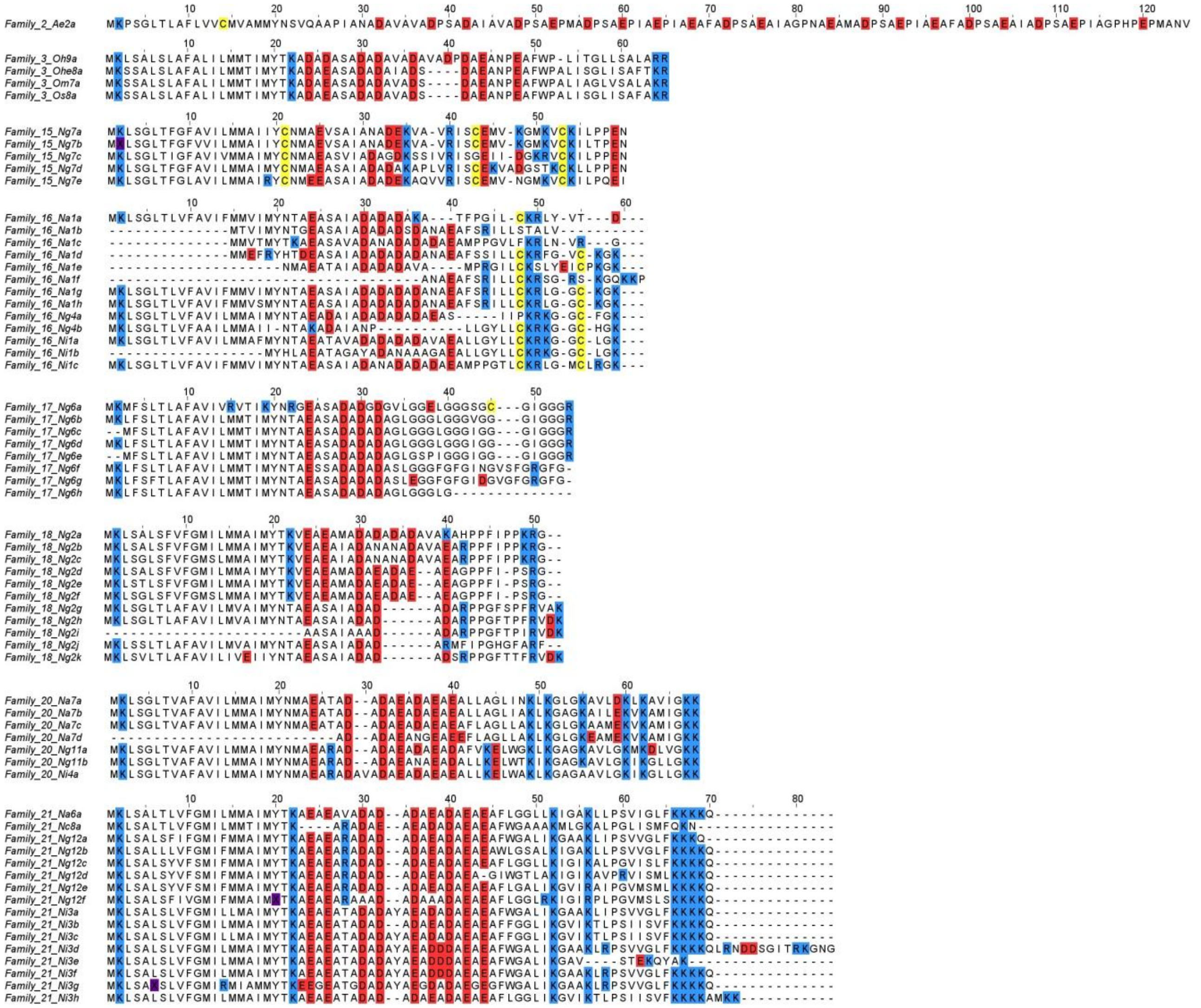

**Figure S12. Alignment of the amino acid sequences of toxin families belonging to the clade of Ponericin-like.** Multiple alignments were generated using the Muscle program in the MEGA-X version 10.1.8 and edited using Jalview version 2.11.2.7. Positively charged residues (lysine and arginine) are highlighted in blue, negatively charged residues (glutamic acid and aspartic acid) are highlighted in red, and cysteine residues are highlighted in yellow.

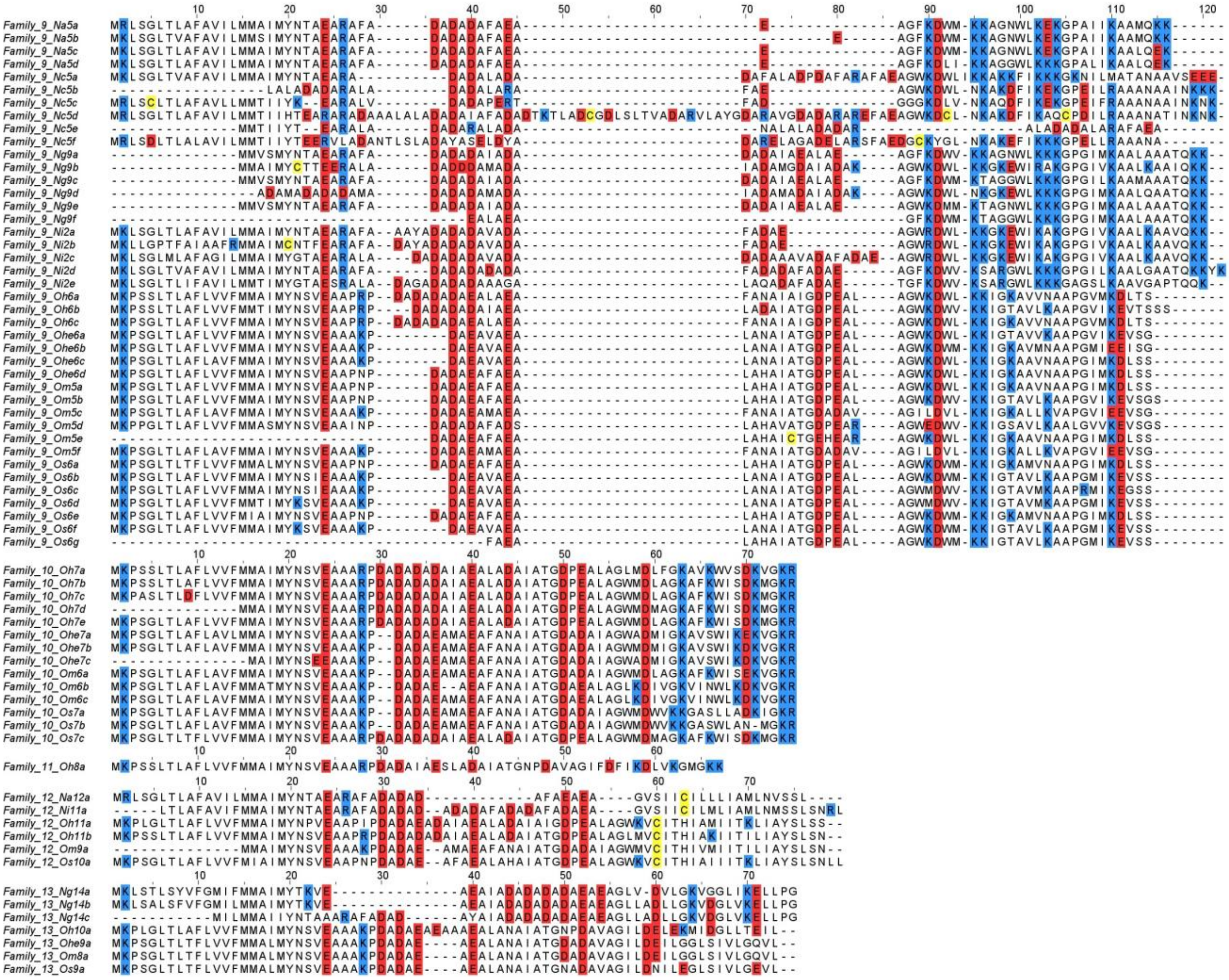

**Figure S13. Alignment of the amino acid sequences of toxin families belonging to the clade of ponericin-like (continued).** Multiple alignments were generated using the Muscle program in the MEGA-X version 10.1.8 and edited using Jalview version 2.11.2.7. Positively charged residues (lysine and arginine) are highlighted in blue, negatively charged residues (glutamic acid and aspartic acid) are highlighted in red, and cysteine residues are highlighted in yellow.

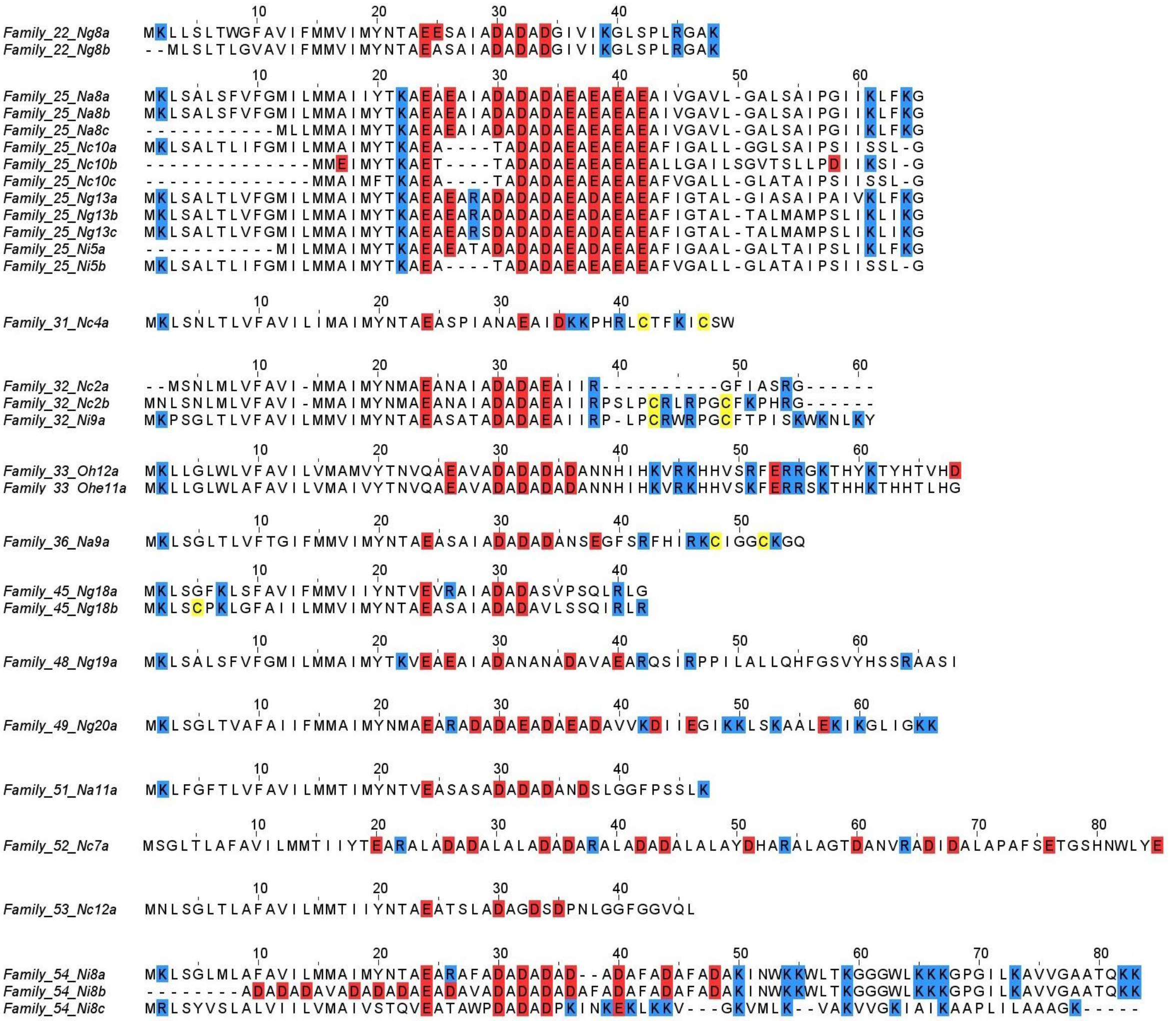


**Figure S14. Alignment of the amino acid sequences of toxin families belonging to the clade of ponericin-like (continued).** Multiple alignments were generated using the Muscle program in the MEGA-X version 10.1.8 and edited using Jalview version 2.11.2.7. Positively charged residues (lysine and arginine) are highlighted in blue, negatively charged residues (glutamic acid and aspartic acid) are highlighted in red, and cysteine residues are highlighted in yellow.

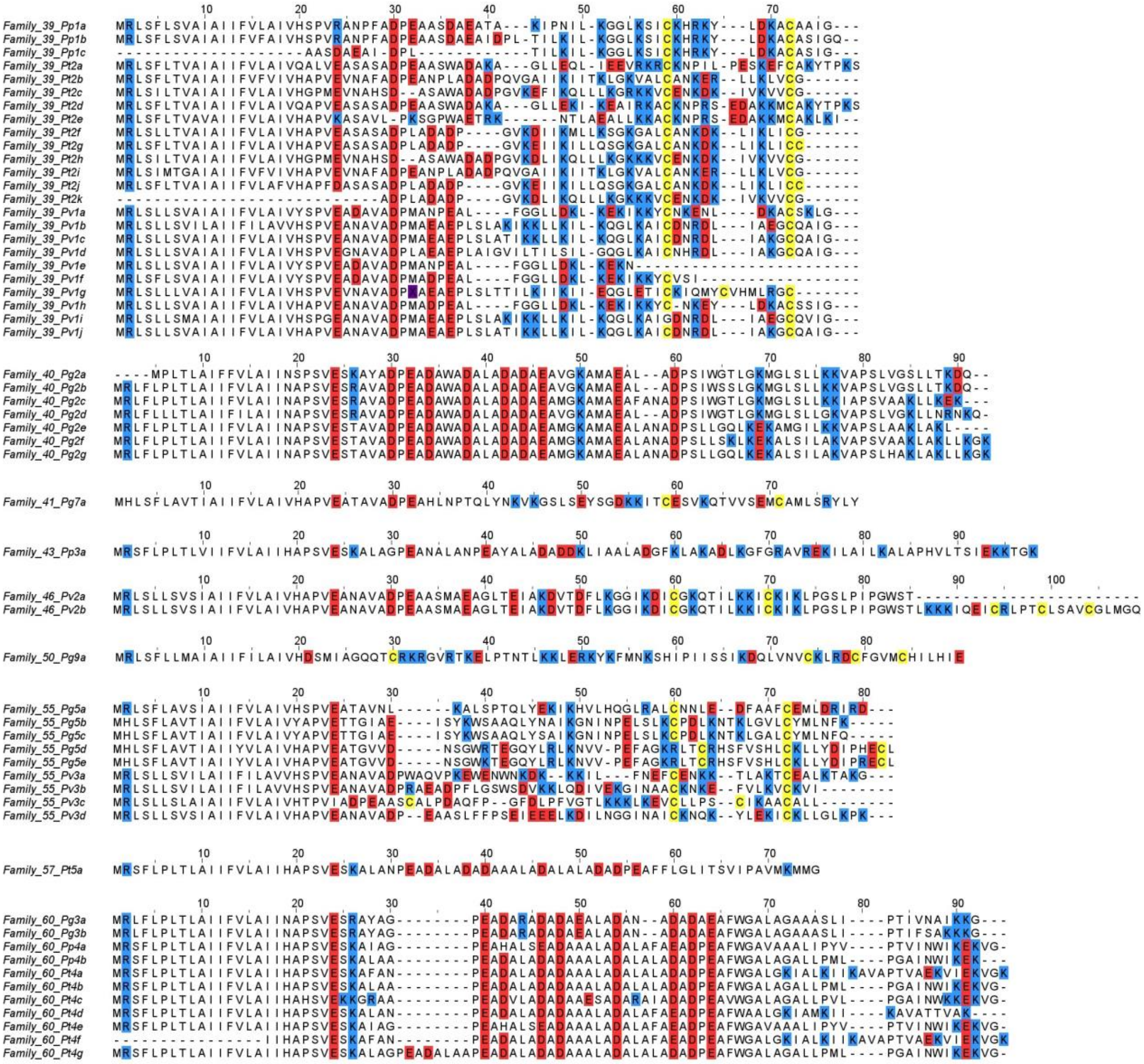

**Figure S15. Alignment of the amino acid sequences of toxin families belonging to the clade of pseudomyrmeciitoxins.** Multiple alignments were generated using the Muscle program in the MEGA-X version 10.1.8 and edited using Jalview version 2.11.2.7. Positively charged residues (lysine and arginine) are highlighted in blue, negatively charged residues (glutamic acid and aspartic acid) are highlighted in red, and cysteine residues are highlighted in yellow.

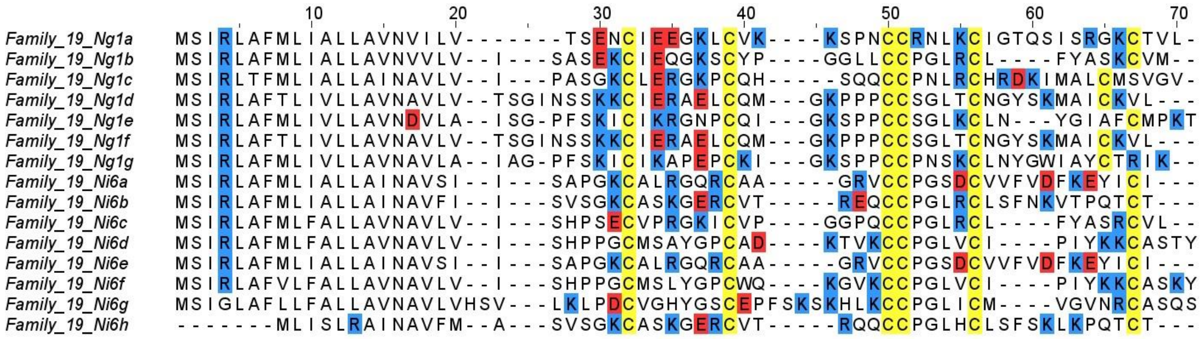


**Figure S16. Alignment of the amino acid sequences of toxin families belonging to the clade of ICK-PONTX.** Multiple alignments were generated using the Muscle program in the MEGA-X version 10.1.8 and edited using Jalview version 2.11.2.7. Positively charged residues (lysine and arginine) are highlighted in blue, negatively charged residues (glutamic acid and aspartic acid) are highlighted in red, and cysteine residues are highlighted in yellow.

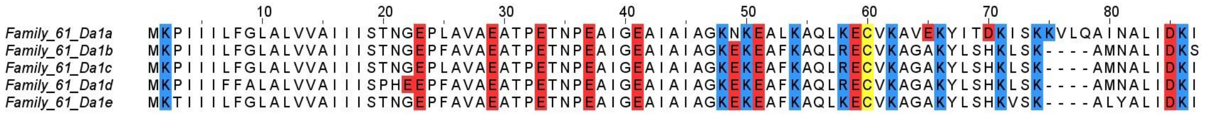


**Figure S17. Alignment of the amino acid sequences of toxin families belonging to the clade of dimeric MYRTX.** Multiple alignments were generated using the Muscle program in the MEGA-X version 10.1.8 and edited using Jalview version 2.11.2.7. Positively charged residues (lysine and arginine) are highlighted in blue, negatively charged residues (glutamic acid and aspartic acid) are highlighted in red, and cysteine residues are highlighted in yellow.

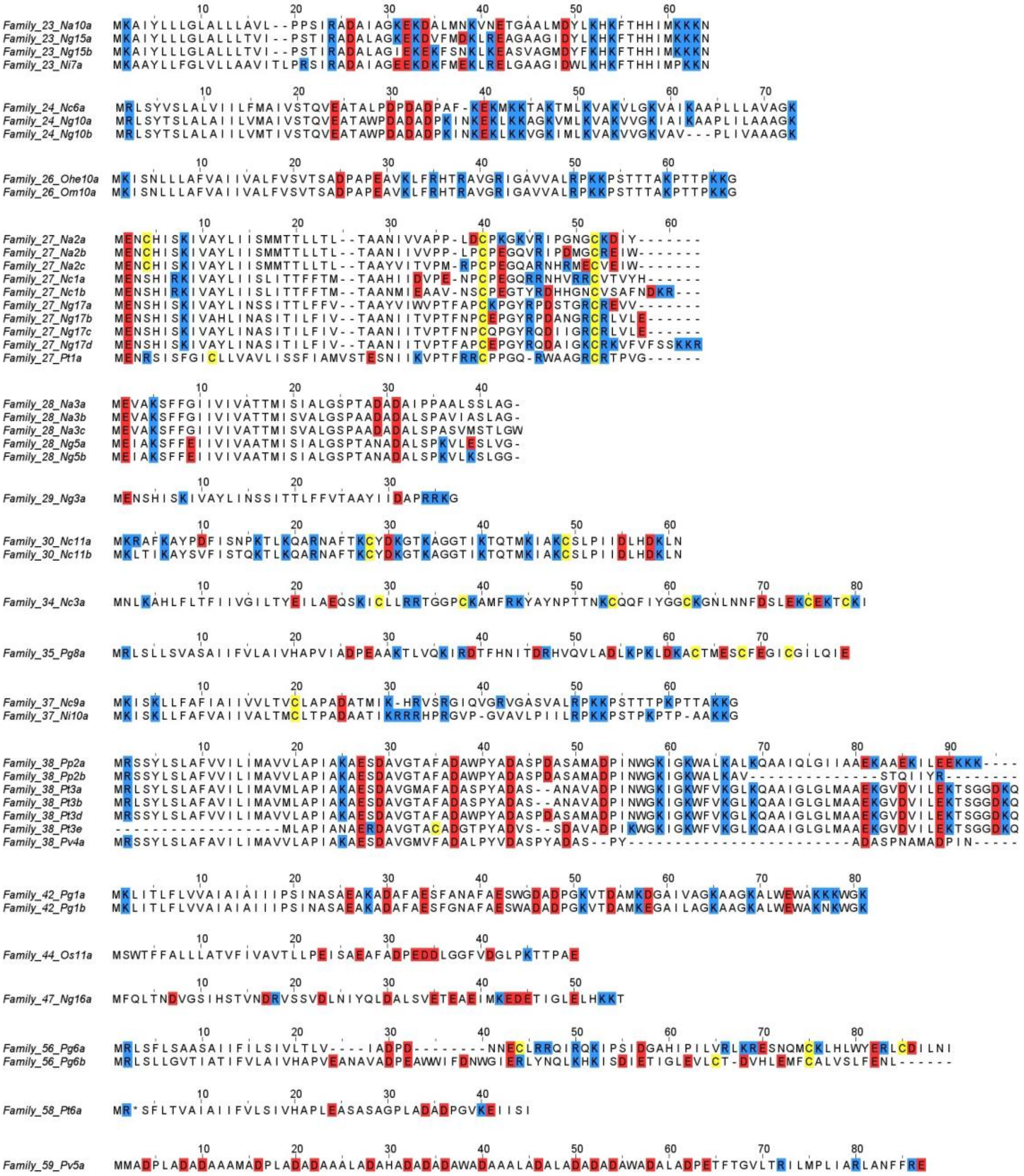

**Figure S18. Alignment of the amino acid sequences of toxin families left as uncategorized.** Multiple alignments were generated using the Muscle program in the MEGA-X version 10.1.8 and edited using Jalview version 2.11.2.7. Positively charged residues (lysine and arginine) are highlighted in blue, negatively charged residues (glutamic acid and aspartic acid) are highlighted in red, and cysteine residues are highlighted in yellow.

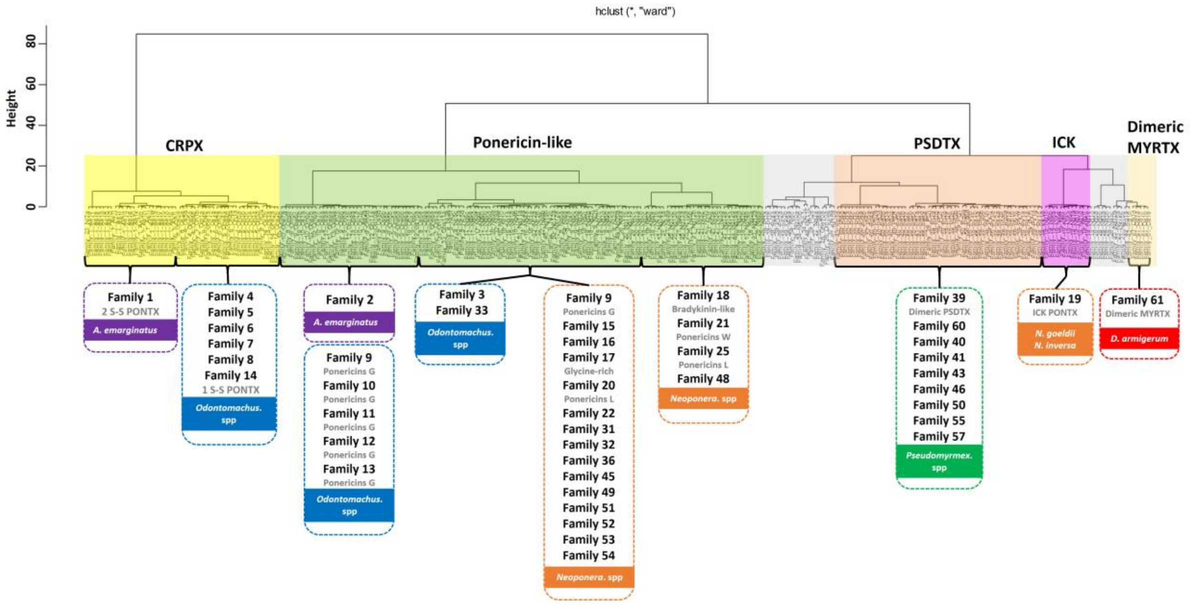


**Figure S19. Classification of peptide toxin transcripts into five precursor clades.** Clustering analysis is based on muscle multiple alignments of all toxin precursor signal regions (determined by SignalP-6.0 https://services.healthtech.dtu.dk/services/SignalP-6.0/), and pairwise distances were calculated using MEGAX software. Final manual verification led us to separate the transcripts into 5 clades, named cysteine-rich poneritoxin (CRPX), ponericin-like, pseudomyrmeciitoxin (PSDTX), ICK and dimeric MYRTX. Gray areas of the cladogram correspond to transcripts with no obvious similarity to others and were left as uncategorized.

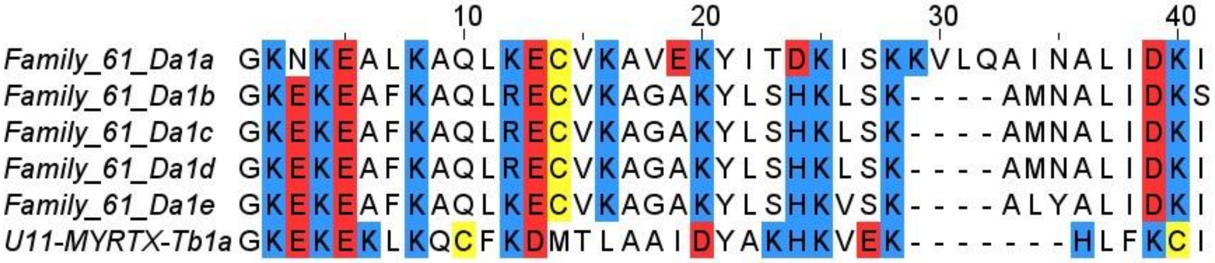


**Figure S20. Alignment of the amino acid sequences of dimeric MYRTX (family 61) and U$_{11}$ peptide from *Tetramorium bicarinatum*.** Multiple alignments were generated using the Muscle program in the MEGA-X version 10.1.8 and edited using Jalview version 2.11.2.7. Positively charged residues (lysine and arginine) are highlighted in blue, negatively charged residues (glutamic acid and aspartic acid) are highlighted in red, and cysteine residues are highlighted in yellow.

**Table S1.** Venom yield and morphological measurements of venom reservoir, sting and mandibles.

| Species | Venom yield (µg) | Venom reservoir | | Sting | | Mandible | |
|---|---|---|---|---|---|---|---|
| | | Volume (nL) | Proportion | Length (mm) | Proportion | Length (mm) | Proportion |
| *Pa. clavata* | 340.4 ± 25.9 | 1785 ± 85 | 6.7 ± 0.5 | 3.6 ± 0.1 | 0.55 ± 0.01 | 4.8 ± 0.1 | 1.0 ± 0.0 |
| *D. armigerum* | 11.5 ± 1.5 | 123 ± 15 | 3.0 ± 0.3 | 1.7 ± 0.0 | 0.49 ± 0.01 | 1.8 ± 0.0 | 1.1 ± 0.0 |
| *A. emarginatus* | 8.0 ± 0.4 | 43 ± 8 | 1.1 ± 0.2 | 1.2 ± 0.0 | 0.35 ± 0.01 | 1.7 ± 0.0 | 0.8 ± 0.0 |
| *O. hastatus* | 106.0 ± 2.5 | 387 ± 38 | 2.9 ± 0.3 | 1.7 ± 0.0 | 0.32 ± 0.00 | 2.2 ± 0.0 | 0.6 ± 0.0 |
| *O. haematodus* | 19.8 ± 2.2 | 129 ± 25 | 3.7 ± 0.6 | 1.2 ± 0.0 | 0.36 ± 0.00 | 1.5 ± 0.0 | 0.6 ± 0.0 |
| *O. scalptus* | 28.0 ± 1.9 | 164 ± 19 | 3.7 ± 0.4 | 1.2 ± 0.0 | 0.33 ± 0.01 | 1.4 ± 0.0 | 0.6 ± 0.0 |
| *O. mayi* | 15.5 ± 1.3 | 96 ± 20 | 3.1 ± 0.5 | 1.3 ± 0.0 | 0.41 ± 0.01 | 1.8 ± 0.0 | 0.6 ± 0.0 |
| *N. goeldii* | 34.7 ± 3.6 | 160 ± 30 | 7.3 ± 1.4 | 1.3 ± 0.0 | 0.47 ± 0.01 | 1.3 ± 0.0 | 0.8 ± 0.0 |
| *N. commutata* | 648.6 ± 52.5 | 4010 ± 368 | 23.2 ± 2.3 | 2.5 ± 0.0 | 0.45 ± 0.00 | 2.8 ± 0.0 | 0.7 ± 0.0 |
| *N. inversa* | 75.7 ± 3.0 | 632 ± 56 | 5.3 ± 0.5 | 2.2 ± 0.0 | 0.45 ± 0.00 | 2.2 ± 0.0 | 0.7 ± 0.0 |
| *N. apicalis* | 81.0 ± 6.0 | 270 ± 30 | 3.4 ± 0.3 | 1.8 ± 0.0 | 0.42 ± 0.00 | 2.0 ± 0.0 | 0.8 ± 0.0 |
| *P. penetrator* | 16.8 ± 1.4 | 79 ± 14 | 20.1 ± 3.3 | 0.9 ± 0.0 | 0.58 ± 0.02 | 0.4 ± 0.0 | 0.4 ± 0.0 |
| *P. viduus* | 20.0 ± 5.4 | 79 ± 12 | 20.5 ± 2.9 | 0.7 ± 0.0 | 0.46 ± 0.01 | 0.4 ± 0.0 | 0.4 ± 0.0 |
| *P. termitarius* | 12.2 ± 1.6 | 53 ± 4 | 6.8 ± 0.6 | 1.0 ± 0.0 | 0.49 ± 0.02 | 0.6 ± 0.0 | 0.5 ± 0.0 |
| *P. gracilis* | 10.5 ± 1.4 | 17 ± 4 | 1.6 ± 0.3 | 0.8 ± 0.0 | 0.38 ± 0.01 | 0.6 ± 0.0 | 0.4 ± 0.0 |

**Table S2.** Effective concentration ($EC_{50}$) of venoms to activate mammalian neuronal cells and nocifensive capacity values.

| Species | $EC_{50}$ (µg/mL) | Nocifensive capacity |
|---|---|---|
| *Pa. clavata* | 3.7 ± 0.2 | 91.9 |
| *D. armigerum* | >100 | <0.01 |
| *A. emarginatus* | 370.1 ± 126.4 | 0.02 |
| *O. hastatus* | 76.1 ± 12.0 | 1.4 |
| *O. haematodus* | 74.8 ± 3.1 | 0.3 |
| *O. scalptus* | 113.8 ± 6.7 | 0.2 |
| *O. mayi* | 68.8 ± 2.9 | 0.2 |
| *N. goeldii* | 26.8 ± 6.1 | 1.3 |
| *N. commutata* | 74.0 ± 6.8 | 8.8 |
| *N. inversa* | 21.0 ± 2.4 | 2.5 |
| *N. apicalis* | 29.6 ± 1.6 | 2.7 |
| *P. penetrator* | 20.3 ± 1.1 | 0.8 |
| *P. viduus* | 138.9 ± 27.0 | 0.1 |
| *P. termitarius* | 53.0 ± 1.3 | 0.2 |
| *P. gracilis* | 97.6 ± 2.8 | 0.1 |

**Table S3.** Paralytic doses ($PD_{50}$) and lethal dose ($LD_{50}$) of venoms against the blowfly *L. caesar*, 1h and 24h after injection.

| Species | $PD_{50}$ (µg/g) | | Paralytic capacity | | $LD_{50}$ (µg/g) | Lethal capacity |
|---|---|---|---|---|---|---|
| | 1h | 24h | 1h | 24h | 24h | 24h |
| *Pa. clavata* | 70.7 ±4.6 | 265.8 ± 14.5 | 4.6 | 1.22 | 414.8 ± 21.4 | 1.73 |
| *D. armigerum* | 5.3 ± 0.2 | 8.2 ± 0.4 | 2.2 | 1.4 | 12.2 ± 0.7 | 0.94 |
| *A. emarginatus* | 7.9 ± 0.2 | 10.0 ± 0.5 | 1.0 | 0.8 | 415.4 ± 18.5 | 0.02 |
| *O. hastatus* | 116.8 ± 8.5 | 188.0 ± 11.0 | 0.9 | 0.6 | 432.0 ± 17.0 | 0,25 |
| *O. haematodus* | 105.6 ± 7.1 | 150.1 ± 12.0 | 0.2 | 0.1 | 374.4 ± 84.9 | 0,05 |
| *O. scalptus* | 179.2 ± 8.8 | 285.3 ± 21.5 | 0.2 | 0.1 | 462.9 ± 20.7 | 0,19 |
| *O. mayi* | 114.3 ± 9.9 | 102.0 ± 7.1 | 0.1 | 0.2 | 504.4 ± 87.5 | 0,03 |
| *N. goeldii* | 47.4 ± 3.4 | 72.8 ± 7.3 | 0.7 | 0.5 | 245.9 ± 35.6 | 0,14 |
| *N. commutata* | 44.4 ± 3.4 | 57.0 ± 2.5 | 14.6 | 11.4 | 265.4 ± 93.6 | 2,44 |
| *N. inversa* | 51.6 ± 5.9 | 65.0 ± 2.8 | 1.5 | 1.2 | 445.5 ± 149.9 | 0,17 |
| *N. apicalis* | 99.6 ± 4.7 | 120.1 ± 7.4 | 0.8 | 0.7 | 265.6 ± 35.0 | 0,30 |
| *P. penetrator* | 7.2 ± 0.4 | 7.0 ± 0.2 | 2.3 | 2.4 | 9.3 ± 0.4 | 1,79 |
| *P. viduus* | 1.2 ± 0.0 | 1.4 ± 0.1 | 16.7 | 14.6 | 2.0 ± 0.1 | 9,85 |
| *P. termitarius* | 12.9 ± 0.9 | 8.1 ± 0.3 | 1.0 | 1.6 | 15.5 ± 1.4 | 0,78 |
| *P. gracilis* | 58.7 ± 1.9 | 47.9 ± 1.8 | 0.2 | 0.2 | 207.9 ± 34.9 | 0.05 |

**Table S4.** Cytotoxicity of crude venoms on S2 *Drosophila* cells based on CCK-8 (metabolism) and LDH (membrane integrity) assays.

| **Species** | **Cytotoxicity $LC_{50}$ (μg/mL)** | |
|---|---|---|
| | **Metabolism** | **Membrane** |
| *Pa. clavata* | 2.72 ± 0.39 | 31.03 ± 9.07 |
| *D. armigerum* | >100 | >100 |
| *A. emarginatus* | >100 | >100 |
| *O. hastatus* | 23.77 ± 4.48 | 36.96 ± 6.32 |
| *O. haematodus* | 17.47 ± 1.59 | 49.02 ± 6.43 |
| *O. scalptus* | 42.82 ± 5.21 | 34.19 ± 8.57 |
| *O. mayi* | 15.87 ± 1.64 | 18.50 ± 2.99 |
| *N. goeldii* | 7.13 ± 0.47 | 7.52 ± 2.17 |
| *N. commutata* | 8.30 ± 0.96 | 5.80 ± 0.67 |
| *N. inversa* | 5.50 ± 0.79 | 5.82 ± 1.51 |
| *N. apicalis* | 5.54 ± 0.90 | 14.88 ± 5.84 |
| *P. penetrator* | 0.05 ± 0.00 | 0.08 ± 0.01 |
| *P. viduus* | 3.40 ± 0.21 | 1.53 ± 0.16 |
| *P. termitarius* | 0.24 ± 0.02 | 0.23 ± 0.02 |
| *P. gracilis* | 4.23 ± 0.26 | 3.18 ± 0.50 |
